# A Template for Translational Bioinformatics: Facilitating Multimodal Data Analyses in Preclinical Models of Neurological Injury

**DOI:** 10.1101/2023.07.17.547582

**Authors:** Hunter A. Gaudio, Viveknarayanan Padmanabhan, William P. Landis, Luiz E. V. Silva, Julia Slovis, Jonathan Starr, M. Katie Weeks, Nicholas J. Widmann, Rodrigo M. Forti, Gerard H. Laurent, Nicolina R. Ranieri, Frank Mi, Rinat E. Degani, Thomas Hallowell, Nile Delso, Hannah Calkins, Christiana Dobrzynski, Sophie Haddad, Shih-Han Kao, Misun Hwang, Lingyun Shi, Wesley B. Baker, Fuchiang Tsui, Ryan W. Morgan, Todd J. Kilbaugh, Tiffany S. Ko

## Abstract

**Background:** Pediatric neurological injury and disease is a critical public health issue due to increasing rates of survival from primary injuries (e.g., cardiac arrest, traumatic brain injury) and a lack of monitoring technologies and therapeutics for the treatment of secondary neurological injury. Translational, preclinical research facilitates the development of solutions to address this growing issue but is hindered by a lack of available data frameworks and standards for the management, processing, and analysis of multimodal data sets.

**Methods:** Here, we present a generalizable data framework that was implemented for large animal research at the Children’s Hospital of Philadelphia to address this technological gap. The presented framework culminates in an interactive dashboard for exploratory analysis and filtered data set download.

**Results:** Compared with existing clinical and preclinical data management solutions, the presented framework accommodates heterogeneous data types (single measure, repeated measures, time series, and imaging), integrates data sets across various experimental models, and facilitates dynamic visualization of integrated data sets. We present a use case of this framework for predictive model development for intra-arrest prediction of cardiopulmonary resuscitation outcome.

**Conclusions:** The described preclinical data framework may serve as a template to aid in data management efforts in other translational research labs that generate heterogeneous data sets and require a dynamic platform that can easily evolve alongside their research.

## Background

Pediatric neurological injury and disease is a critical public health issue (1–3) but is challenging to characterize due to relatively small patient populations and wide heterogeneity compared to adult clinical populations. Maturation of the developing brain (e.g., synaptogenesis, myelination, and cortical development) leads to rapid changes in cerebral hemodynamics and metabolism in children (4), (5), causing management of pediatric neurological injury and associated risk factors to be distinct from adults. Children who suffer primary traumatic brain injury (TBI) or cardiovascular and respiratory injury experience secondary neurological injury that is often uncharacterized and untreated and is associated with worse functional outcomes (6–9). Recent advancements in critical care technology and treatment have significantly improved survival following primary neurological injury (10) and in other populations in which secondary neurological injury is common (e.g., congenital heart disease) (11). The development of interventional strategies to address persistently elevated rates of adverse neurological outcomes in this growing population is urgently needed (12). A deeper understanding of the mechanisms that drive primary and secondary neurological injury is required to develop targeted monitoring technologies and precision therapeutics to help improve outcomes.

The diverse presentation of pediatric disease leads to wide patient-to-patient heterogeneity; this clinical heterogeneity confounds characterization of neurological injury mechanisms and necessitates large sample sizes for meaningful stratification. Preclinical injury and disease models provide the opportunity to characterize neurological injury mechanisms in a controlled and repeatable environment. In particular, large animal swine models are uniquely suited to model pediatric (neonatal to adolescent) neurological injury due to functional, morphological, and maturational similarities with the human brain (13–17). Of major importance, the size of the pig brain is ideally suited for clinically relevant, biomechanical modeling of human head trauma (18–21), including both focal and inertial injuries to gray and white matter, as well as the development of novel optical neuromonitoring techniques, where distance from the surface of the skin to the brain and optical properties of the scalp tissue and skull are easily translated to human clinical populations (22), (23). Swine have been recognized as a common preclinical model species for TBI by the Federal Interagency Traumatic Brain Injury Research (FITBIR) organization (https://fitbir.nih.gov/).

In addition to high-fidelity animal models, the preclinical environment permits the collection of a myriad of experimental parameters, such as pathology, hemodynamics, omics data sets, and other downstream, derived parameters, to characterize neurological injury and associated mechanisms. Data-driven analysis of these physiological data sets enables novel identification of mechanisms of neurological injury and potential therapeutic pathways to advance clinical medicine. However, integration of these diverse and complex data sources poses a significant logistical hurdle, and when done non-uniformly, hinders data reuse which may obscure valuable physiologic insights.

Current clinical data management systems are ill-equipped for storage and analysis of high-resolution time series data (24). We refer to sampling rates of ≥ 10 Hz as high-resolution data due to the ability to resolve time series fluctuations at the frequency of the cardiac cycle. This results in the collection of “convenience samples” where patient sampling may not be representative and is non-uniformly catalogued. Furthermore, clinical data management systems are not compatible with custom, investigational monitoring systems that may yield significant future utility and are under development in high-fidelity large animal models (25). Non-uniform data cataloguing, temporal synchronization, and health data privacy considerations pose significant obstacles to cross-sectional indexing of clinical events that may lead to neurological injury, such as cardiac arrest (26), (27). Barriers to the integration and standardization of multimodal data for downstream, cross-sectional analyses lead to significant data loss and limit scientific discovery (28).

The use of standardized data frameworks for preclinical research remains rare yet holds significant potential for maintaining rigor in prospective analyses and simultaneously facilitating collaborative, retrospective inquiry. By examining historical data sets, the need for additional exploratory animal experiments may be reduced. Preclinical data sets have traditionally been shared via publication, as averaged, down sampled aggregates, and do not benefit from common data element (CDE) naming conventions for interoperability (29). To address this gap, we developed a novel, generalizable preclinical data framework for the integration and visualization of multimodal data sets across heterogeneous experimental models of pediatric neurological injury.

In this work, we describe the data sources and data types collected within our institution, and the processing pipeline that was developed for integration, analysis, extract, transform, load (ETL), display and export. Leveraging our laboratories’ expertise in a variety of large animal models, this framework features compatibility with data derived from our cardiac arrest, TBI, cardiopulmonary bypass, and extracorporeal models among others. Furthermore, we present a use case for our framework that enables the discovery of novel predictors of resuscitation success and neurological injury in a preclinical cardiac arrest model. This novel data framework serves as a template to increase experimental data reusability through data stewardship.

## Methods

The Resuscitation Science Center (RSC) of Emphasis at the Children’s Hospital of Philadelphia (CHOP) Research Institute conducts preclinical research, assessing risk factors for, and testing novel therapeutics to treat neurological injury in animal models. All animal procedures were approved by the CHOP Institutional Animal Care and Use Committee (IACUC) and were conducted in strict accordance with the NIH Guide for the Care and Use of Laboratory Animals. From 2017-2022, 747 animal studies were conducted across a range of translational neurological injury models (53.0% cardiac arrest, 16.9% cardiopulmonary bypass, 15.8% traumatic brain injury, 6.6% carbon monoxide toxicity, 5.0% hydrocephalus, 2.7% neonatal hypoxia). Animals ranged in age from newborns to 120 days, corresponding to a weight range of ∼1 to ∼40 kg. From these experiments, raw data types collected include single measure, repeated measures, time series, and imaging. To facilitate analyses utilizing these heterogeneous data types, the development of a custom data framework was needed. Design considerations included rapidity of processing, near-full automation, and adaptability to accommodate potential future data modalities. The developed data framework encompasses specific data collection tools and systems that support investigational monitoring devices, data processing pipelines that feature data quality checkpoints, tailored ETL methodology for data warehousing, and a visualization platform that permits dynamic data filtering, visualization, and export.

To clarify the terminology we will use, “data type” refers to the category of data being collected (“single measure”, “repeated measures”, “time series”, or “imaging”). “Data modality” refers to groups of recorded “data elements”. If the data modality is “Arterial Blood Gas”, a data element is a measured variable within the modality (e.g., “pH”). The “experimental model” refers to the general injury or therapeutic model. The “cohort” refers to a specific study within that experimental model. Last, the “subject” refers to an individual experiment, classified by the subject’s unique identifier, which combines the experimental date in the format “YYMMDD” and the number assigned to the animal by the vendor (e.g., “210813_1745”). These naming conventions facilitate multileveled organization and comparative analysis.

### File Structure

A standardized, hierarchical file structure provided organization of data by experimental model, cohort, collection instrument, and processing stage (Fig. 1). Our team collaborated with the CHOP Research Institute Arcus Library Science Team (H.C., C.D.) to design an optimized file structure.

**Fig. 1.**
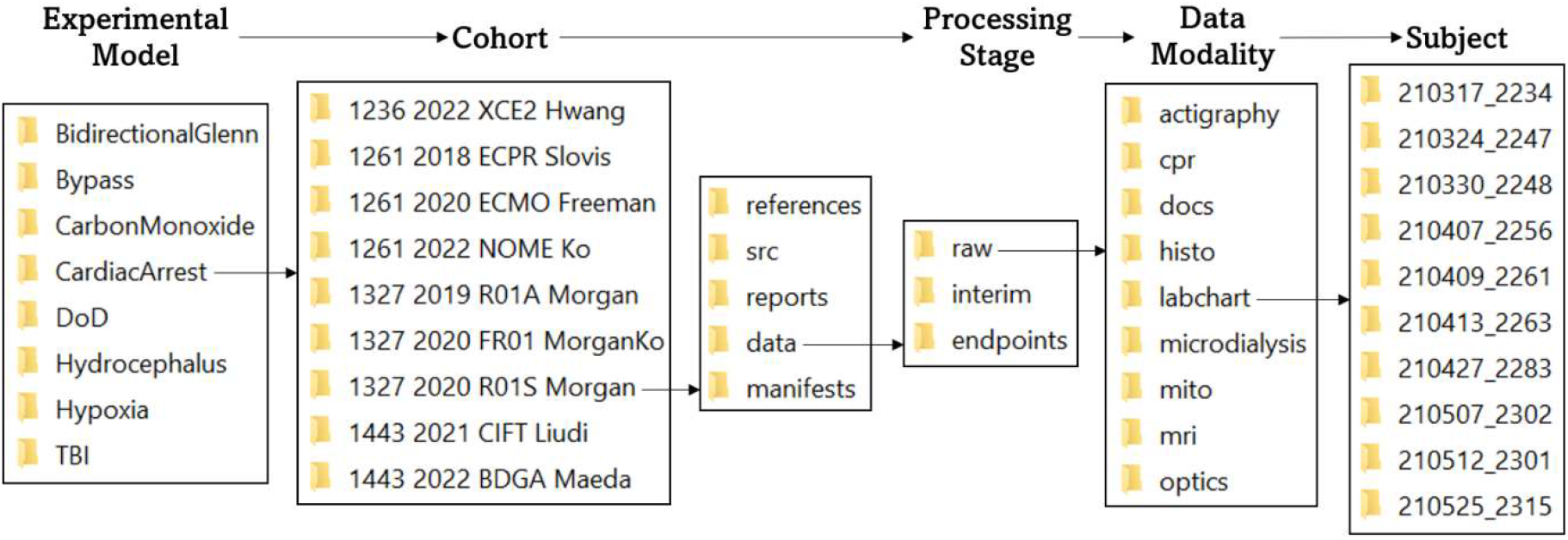
File Structure Diagram: This diagram demonstrates hierarchical organization of the file structure by classifier (experimental model, cohort, subject), processing stage, and data modality. This nested structure facilitates the storage of metadata within the file’s path name, which is more efficient than accessing legend files for downstream data integration and requires no manual updating or quality review. The organization of this file structure allows data to be queried by classifier, processing stage, and data modality, and provides space for the storage of cohort-specific documentation and analysis files.

The file structure described below is hierarchically organized using the classifiers “experimental model”, “cohort”, and “subject”. Each experimental model is identified by a unique name that reflects the model (e.g., “CardiacArrest” for cardiac arrest, “TBI” for traumatic brain injury). To date, twenty-two unique cohorts are contained within the “CardiacArrest” directory. The syntax for cohort directory naming is a combination of the CHOP IACUC protocol number, the year of the first experiment, a unique four-character alphanumeric cohort identifier, and the principal investigator’s last name (e.g., 1261 2022 NOME Ko). Many data modalities are shared by cohorts within the same experimental model. Protocolized data type storage across different experimental models and cohorts enables comparative, cross-sectional data element analyses. Within each cohort directory, data at various stages of processing are contained within a “data” folder. In addition to housing raw experimental data types, the file structure within this folder features designated directories for downstream data processing that facilitate workflow tracking to monitor the processing status of each data modality. Specifically, “raw”, “interim”, and “endpoints” folders indicate the processing status of each data type, and ensure that only processed, quality-reviewed data are integrated and transferred to various locations for sharing, visualization, and further analysis.

Alongside the “data” folder, each cohort directory also contains a “manifest”, “src”, “reports”, and “references” folder. The “manifest” folder contains cohort-wide generated sheets, summarizing the data collection and processing status of each data modality (see “Manifest Generation”). The “src” folder contains code and other supporting material for cohort-specific analyses and the “reports” folder contains the results generated by these analyses. Lastly, the “references” folder contains reference material related to the cohort’s experimental design.

### Data Types & Data Collection

Diverse, multimodal, preclinical data types are collected during each experiment. Data that are acquired once per subject are referred to as “single measure” data. “Repeated measures” data are acquired at multiple timepoints per subject. These timepoints may be predefined or change from subject to subject. “Time series” data are sequences of data indexed at successive, equally spaced points in time. A distinction between repeated measures data and time series data is that intervals between repeated measures data are often irregular and at the time scale of hours to days. Finally, the “imaging” data type describes spatial data (greater than one dimension) that may yield derived data types that are single measure, repeated measures, or time series. Standardized collection instruments were tailored to support specific data types and collection environments and are further described below.

### Single Measure Data

For each experiment, single measure data including demographic information about the subject, the monitoring and data collection techniques used during the experiment, any medications or interventions administered to the subject, and qualitative and quantitative characteristics of the experimental protocol are collected. Examples of collected single measure data include species, sex, date of birth, experimental group, and types of anesthesia administered. The collection of these data allows full understanding of all aspects of an experiment, which is essential to the analysis and interpretation of collected physiological data. Single measure data may also be associated with a timestamp; this includes experimental outcomes data that may occur at different experimental timepoints depending upon survival duration. Timestamped, single measure data elements implemented within the platform include quantitative assessments of collected tissue samples (e.g., density of microglia staining in cortical tissue, brain tissue water fraction). Timepoint collection permits cross-sectional analysis of the association of the outcome metric with time and experimental period. The collection tools and methodology of use for single measure data are described below.

### Case Report Forms (CRFs)

CRFs are used in clinical trial research to protocolize the collection of patient data and facilitate reporting of these data to sponsoring institutions (30). We have translated this data collection practice to preclinical research to standardize the collection of data types, including experimental protocol characteristics and physiological measurements during experiments. For each subject, three unique CRFs are used to record qualitative and quantitative information during the experiment: the “Anesthesia Record”, “Code Leader”, and the “Neuro-Optical Monitoring CRF”.

The “Anesthesia Record” records all experimental, therapeutic, and surgical interventions as well as administration quantities of anesthetic and other drugs. Additionally, hemodynamic and ventilation parameters are manually recorded every 5-10 minutes to ensure the wellbeing of the animal. Sharing of this document with CHOP’s Department of Veterinary Resources is mandated by the United States Department of Agriculture (USDA) to ensure humane treatment of each subject.

The “Code Leader” document is used to record clinically relevant information during the injury and/or therapy portion of the experimental protocol. For instance, the Code Leader document is used during the resuscitation portion of an experiment designed to compare the efficacy of different resuscitation techniques or technologies. During this portion of the experiment, the Code Leader document is used to record the timing of vasopressor administration and defibrillation attempts, as well as “clinical” observations.

Finally, the “Neuro-Optical Monitoring CRF” is used for recording observations with precise timestamping, focusing on experimental and physiological parameters that may affect cerebral perfusion or neuro-optical monitoring signal quality. All three documents are scanned and uploaded into the subject’s folder within the cohort’s “docs” folder at the conclusion of the experiment.

Experimentally relevant single measure and repeated measures data are then manually entered into a customized electronic data capture software.

### Electronic Data Capture Software

*REDCap* (https://projectredcap.org/software/) is a customizable, browser-based, metadata-driven electronic data capture software commonly used in translational research for the collection of single measure and repeated measures data types (31), (32). *REDCap* “Arms” are constructs that allow study events to be grouped into a sequence. In our implementation, each “Arm” in the *REDCap* database represents an experimental cohort and is named using the cohort naming convention (see “File Structure”). “Instruments” are the individual data capture forms that prompt the user for data entry. Our *REDCap* instruments are used to collect single measure data including subject characteristics (e.g., “Demographics”, “Monitoring”, “Tissue Collection”) as well as data specific to experimental periods (e.g., “Baseline”, “Asphyxia”, “Acute Survival”). Subject characteristics include data collection checklists that are used to track the collection and usability of various measurement modalities (see “Manifest Generation”). When a new cohort is implemented in the *REDCap* database, only the instruments relevant to that experimental model are selected, with the ability for cohort-level customization. This reduces empty fields and creates a streamlined data entry process.

### Biological Sample Collection Characteristics

Single measure data related to stored biological samples were input into *Freezerworks (Dataworks Development, Mountlake Terrace, WA, USA)* software. This includes information about the sample, the collection protocol, and its location within the center’s freezer system. The *Freezerworks* database was designed to categorize samples by cohort and experimental model for compatibility with other in-house data collection systems and processing pipelines.

### Assessment of Mitochondrial Respiration

Mitochondrial respiration is a metric of the oxidative respiratory capacity of the mitochondria and is assessed at a single experimental timepoint from fresh tissue sampled at euthanasia. A significant reduction in respiratory capacity has been associated with adverse functional neurological outcomes (33). High-resolution respirometry is used to measure mitochondrial respiration in brain tissue homogenate in an *Oxygraph-2k (Oroboros Instruments, Innsbruck, Austria)*. Additions of complex-specific substrates and inhibitors allow for the assessment of the integrated respiratory capacities of oxidative phosphorylation (OXPHOS) and the electron transport system (ETS). Key data points of interest include the oxidative phosphorylation capacity of complex I (OXPHOS_CI) alone and the convergent capacity of complexes I and II (OXPHOS_CI+CII), assessment of the leakiness of the mitochondrial membrane, maximal convergent non-phosphorylating capacity of complexes I and II through the electron transport system (ETS_CI+CII), ETS capacity through complex II alone (ETS_CII), and ETS capacity through complex IV alone (ETS_CIV). To normalize quantitative respiration values for mitochondrial content, samples are also assessed for citrate synthase (CS) activity using a commercially available kit, *Citrate Synthase Assay Kit CS0720 (Sigma-Aldrich, St. Louis, MO, USA)*. Raw respirometry data files (.*dld*) produced by the *Oxygraphy-2k* must be viewed in *Oroboros DatLab* software and then manually exported as a numerical data table (.*cs*v) to a designated “mito” processing directory in the “raw” data folder, where they are later queried by a custom *MATLAB (MathWorks, Natick, MA, USA)* script for automated processing of respirometry values.

### Histopathological Quantification

Histopathological quantification of collected tissue samples yields single measure data that will be subsequently discussed in the “Imaging” section.

### Repeated Measures Data

Repeated measures data are captured in “repeated instrument” forms within *REDCap*, as well as in *.csv* data tables generated from analyses of biological samples and bedside imaging. Collection tools and methodology of use for repeated measures data are described below.

### Electronic Data Capture Software

The custom *REDCap* database stores repeated measures data (e.g., “Blood Gasses”, “Cardiac Outputs”) utilizing “repeating instruments” to capture all instances of collection, without having to first specify the number of instances. These “repeating instruments” are experimentally timestamped and are later aligned with time series data during the “interim” processing stage to provide additional physiological context.

### Cerebral Microdialysis Quantification

Interstitial fluid from cerebral tissue is invasively collected using a *70 Microdialysis Catheter (M Dialysis Inc., Stockholm, Sweden)*. The collected dialysate fluid contains metabolites that may be used to characterize neurological injury. Cerebral microdialysis samples are analyzed on the *ISCUSflex Microdialysis Analyzer (M Dialysis Inc. Stockholm, Sweden)*. The reagent kit used analyzes the samples for levels of glucose (mg/dL), lactate (mM), pyruvate (µM), and glycerol (µM). Data from the analyzer is exported via a universal serial bus (USB) hard drive and viewed using *ICUpilot (M Dialysis Inc., Stockholm, Sweden)* software. The data is then exported from the *ICUpilot* software as a *Microsoft Excel (Microsoft Corporation, Redmond, WA, USA)* spreadsheet and later queried for experimental time-alignment in the “interim” processing stage.

### Venous and Arterial Blood Gas Analysis

Blood gas analysis measures the amounts of dissolved gases (oxygen and carbon dioxide), acidity (pH), electrolyte and metabolite concentrations, and co-oximetry from a collected blood sample. Blood gas values from venous and arterial whole blood samples are measured by a *GEM Premier 5000 (Werfen, Barcelona, Spain)* blood gas analyzer at numerous predefined timepoints for each cohort, manually recorded in the “Code Leader” CRF, and inputted into the repeating *REDCap* instrument, “Blood Gasses”. Each measurement set of blood gasses is entered into a new instance of the repeating instrument within the *REDCap* entry for the subject and is thus able to be individually queried during the “interim” processing stage.

### Cardiac Output Measurement

Cardiac output is the product of heart rate and stroke volume, measured in liters per minute (L/min). Values are measured by an *Agilent V24CT Patient Monitor (Agilent Technologies, Santa Clara, CA, USA)* at numerous predefined timepoints, manually recorded in the “Code Leader” CRF, and inputted into the repeating *REDCap* instrument, “Cardiac Outputs”. Each measurement of cardiac output is entered into a new instance of the repeating instrument within the *REDCap* entry for the subject. These data later undergo subject-specific integration during the “interim” processing stage.

### Protein and Metabolite Quantification

Mass spectrometry of blood serum or plasma samples collected throughout an experiment yields protein and metabolite data that grant a window into the dynamic environment of the proteome and metabolome, respectively. From these data, biomarkers (either protein products or metabolites) of interest may be analyzed over the course of recovery, or as a point of comparison for different groups (34), (35). Plasma biomarkers are analyzed through the Penn Medicine Human Immunology Core (HIC) using *Simoa® (Quanterix Corp., Billerica, MA, USA)* assay kits. The HIC compiles assay plate results into an *Excel* spreadsheet. A custom *MATLAB* script extracts biomarker data from corresponding spreadsheets, renames data points, and saves these data in *.csv* files. The *.csv* files are then imported into *R (R Foundation for Statistical Computing, Vienna, Austria),* where statistical analysis is performed. Statistical results are saved as *.csv* files in the subject’s folder within the “interim” directory, along with individual *.png* files for the generated figures. These data later undergo subject-specific integration during the “interim” processing stage.

### Imaging

In many cases, imaging data are collected at numerous experimental timepoints and thus are also deemed a repeated measures data type. The unique types of imaging data collected will be elaborated on in the “Imaging” data type section due to heterogeneity and complexity.

### Time Series Data

Several physiological time series data elements are collected throughout each experiment. The collection frequency and period vary by collection instrument. Collection hardware and software for time series data are described below.

### Hemodynamic and Other Physiological Waveforms

*PowerLab* (*ADInstruments, Sydney, Australia*) hardware modules collect physiological time series data at 1000 Hz. For compatibility with *PowerLab*, the physiological monitors used must have the capability to export their waveforms as analog output voltage signals. *LabChart* (*ADInstruments, Sydney, Australia*) physiological data analysis software centralizes these waveforms and allows the export of all physiological data in one *LabChart* data file (.*adicht*) on the same synchronized time axis. Examples of these physiological waveforms include capnography, rectal temperature, and invasively measured arterial blood pressure. Additionally, cyclic measurements may be derived from waveforms with cyclic features associated with the cardiac or respiratory cycle. Examples of this include the diastolic and systolic pressure calculations from the aortic pressure (AoP), right atrial pressure (RAP), and pulmonary artery pressure (PAP) waveforms and the end-tidal carbon dioxide (EtCO2) measurement from the capnography waveform. Coronary perfusion pressure (CoPP) is calculated as the difference between the diastolic AoP and diastolic RAP values. These calculated waveforms are then saved and exported distinctly from their derivative waveforms. Calculated waveforms combining two different raw waveforms (e.g., CoPP) are uniquely possible in the preclinical setting and have informed numerous studies on the physiologic effects of CPR (33,36–39).

The *LabChart.adicht* file is read and maintained in its native form for some analyses, while the data are divided by experimental periods and downsampled for other analyses. During downsampling, each variable is averaged into 15-second epochs to strike a balance between information density and ease of data export and visualization. These downsampled time series data are used for low-resolution waveform analysis during the “raw” processing stage.

### Cardiopulmonary Resuscitation (CPR) Chest Compression Mechanics

During CPR, a *ZOLL R-Series Advanced Life Support Automated External Defibrillator (ZOLL Medical Corporation, Chelmsford, MA, USA)* with CPR quality monitoring capabilities is used to record compression mechanics data such as depth, rate, and release velocity. These time series data are exported from the hardware as a .*ful* file and are viewed in and exported from *RescueNet Code Review (Zoll Medical Corporation)* software as an *.xml* document. These time series data are time-aligned with the “CPR” experimental period during the “interim” processing stage.

### Actigraphy

The activity level of the subject is measured using an attached accelerometer. This provides an assessment of normal activity as an indicator of neurological impairment following injury. Accelerometer data are collected using a *USB Accelerometer X16-1Eand (Gulf Coast Data Concepts, LLC, Waveland, MS, USA)* and are logged on a battery-powered unit that holds 2-3 days of accelerometer data. These data are then downloaded as a *.txt* file for further “raw” processing and subject-level integration during the “interim” processing stage.

### Electroencephalography

A two-channel differential electroencephalography (EEG) montage is used to record EEG waveforms between C3-P3 locations on the left hemisphere and C4-P4 locations on the right hemisphere, according to the 10-20 system (40). EEG waveforms are captured at a sampling rate of 256 Hz on a clinical bedside monitoring device, *CNS-100R (Moberg Research, Ambler, PA, USA)*. This device is manually time-synchronized during each recording session. Recording sessions are exported and translated from their proprietary data format into a .*mat* data file for import into *MATLAB* for further processing and alignment with other time series data during the “interim” processing stage.

### Neurometabolic Optical Monitoring (NOM)

Continuous, non-invasive, diffuse optical monitoring of cerebral hemodynamics is conducted using an investigational research instrument that has been previously described (41). This research instrument combines the techniques of frequency-domain and broadband diffuse optical spectroscopy (FD-DOS and bDOS, respectively) and diffuse correlation spectroscopy (DCS) via a non-invasive optical sensor that is secured to the animal’s forehead. FD-DOS quantifies cerebral tissue optical scattering and absorption properties from which the tissue concentration of oxy-, deoxy-, and total hemoglobin and cerebral tissue oxygen saturation (StO2) may be derived at a sampling rate of 10 Hz. Using bDOS in conjunction with FD-DOS significantly increases spectral sampling and permits additional quantification of changes in the tissue concentration of water, lipids, and cytochrome c oxidase. bDOS data may be acquired at 1 Hz and is temporally interleaved with FD-DOS and DCS. DCS measures relative changes in cerebral blood flow at a sampling rate of 20 Hz.

Raw NOM waveforms include measured amplitude and phase with respect to modulation frequency, source wavelength and source-detector separation from FD-DOS, absorption spectra from bDOS, and normalized temporal intensity autocorrelation curves and intensities for each detector from DCS. These raw data are imported into *MATLAB* for further processing during the “raw” processing stage. During “raw” processing, theoretical models of light propagation in tissue are used to numerically derive the value of physiological parameters. NOM-derived physiological parameters are integrated with other physiological time series during the “interim” processing stage. It is important to note that derived physiologic parameters vary based on model selection; this poses a challenge to multi-institution data aggregation. Data standardization and optimal model selection are ongoing areas of research in diffuse optics.

### Imaging Data

Imaging may yield data that are single measure, repeated measures, or time series. We discuss this data type separately due to its unique storage and processing considerations. Imaging data vary in size and type based on modality, dimensionality, number of frames (single image vs. video), and resolution. Raw imaging data are stored in various databases and present unique accessibility challenges compared to the other collected data types. Collection tools, storage platforms, and methodology of use for imaging data are described below.

### Magnetic Resonance Imaging (MRI)

MRI data are acquired using a *3T Tim Trio (Siemens Medical Solutions USA, Malvern, PA, USA)*. During each MRI session, several individual scan sequences are performed and stored on the scanner under the subject’s identifier. Sequences include anatomical (e.g., T1, T2) and functional imaging (e.g., SWI, DTI, MRS, ASL, BOLD) (34), (35). Scan data are stored as *.dicom* files. The scanner is outfitted with a *Flywheel (Flywheel Software, San Francisco, CA, USA)* “reaper” that moves data from the scanner to *Flywheel*.

*Flywheel* is a biomedical research data management platform for imaging and associated data. The scanner reaper is configured in a push setup, such that when imaging sessions are completed, the reaper collects the imaging files and pushes them to a *Flywheel* receiver that distributes them to specific folders in *Flywheel.* Once in *Flywheel*, .*dicom* files are viewable with *Flywheel’s* web-based image viewer platform. All files are downloaded for “raw” processing and further analysis. Categorical and numerical parameters derived from neuroimaging include presence, location, and size of neuroimaging abnormalities and the location and value of spectroscopic analytes. These values are integrated by cohort during the “interim” processing stage. Standardization of derived parameters remains ongoing.

### Histology

Numerous tissue samples are collected at euthanasia for histologic analysis of cellular injury and functional impairment (34), (36). Sampled organs include the brain, heart, and kidney. Tissue samples are formalin-fixed, paraffin-embedded, and sectioned at 5-10 µm for chromogenic and immunofluorescent staining. Images of stained tissue sections are taken on a *DMi8 (Leica Microsystems, Wetzlar, Germany)* microscope using a *FLEXCAM C1 (Leica Microsystems, Wetzlar, Germany)* color camera at 20x magnification. Images are then exported as .*tiff* files into *ImageJ* (https://imagej.nih.gov/ij/), where they are deconvoluted using the “H DAB” function. Thresholding is applied only to the “Colour 2” output to measure positive antibody expression. The preset “Measure” function is used to obtain the area, mean, minimum pixel value, and maximum pixel value for each image. Images and associated .*csv* files containing quantitative data are stored in a custom histology file architecture where they can be queried for subject-level integration during the “interim” processing stage.

### Contrast-Enhanced Ultrasound (CEUS)

CEUS data are acquired using both a clinical ultrasound scanner (*Acuson Sequoia*; *Siemens Medical Solutions USA, Malvern, PA, USA*) and a research ultrasound scanner (*Vantage 256; Verasonics Inc., Kirkland, WA, USA*) (42–44). For the clinical ultrasound scanner, video clips or “cineloops” of clinically relevant timepoints during contrast perfusion are manually saved by the sonographer to the scanner’s internal storage as .*dicom* files. After the study, the .*dicom* files are copied to an external USB hard drive. For the research ultrasound scanner, scanning with the probe populates the local machine memory with radio frequency (RF) signal data. The data is asynchronously transferred to a host controller computer via direct memory access according to programmatic requests from custom software written in-house. Once on the host controller, the data is temporarily stored within internal random-access memory (160 GB capacity), and then saved in parallel threads to an internal hard drive storage (28 TB capacity). The RF data can then be processed into images for further visual or computational analysis during “raw” processing. Derived quantitative values are integrated by subject during the “interim” processing stage.

### Data Processing

A custom data integration and processing pipeline was developed for time synchronization, quality-review, and uniform export of collected data. A flow chart demonstrating the functionality of the three data processing stages (“raw”, “interim”, and “endpoints”) can be found in Fig. 2. The pipeline is executed on a subject-by-subject basis during the “raw” and “interim” processing stages, while “endpoints” processing is conducted at a cohort-level. The “raw”, “interim”, and “endpoints” stages populate the corresponding folders of the “data” directory in the file structure (see “File Structure”). This framework provides a standardized, modular system to incorporate novel data types that are a crucial aspect of preclinical research.

**Fig. 2.**
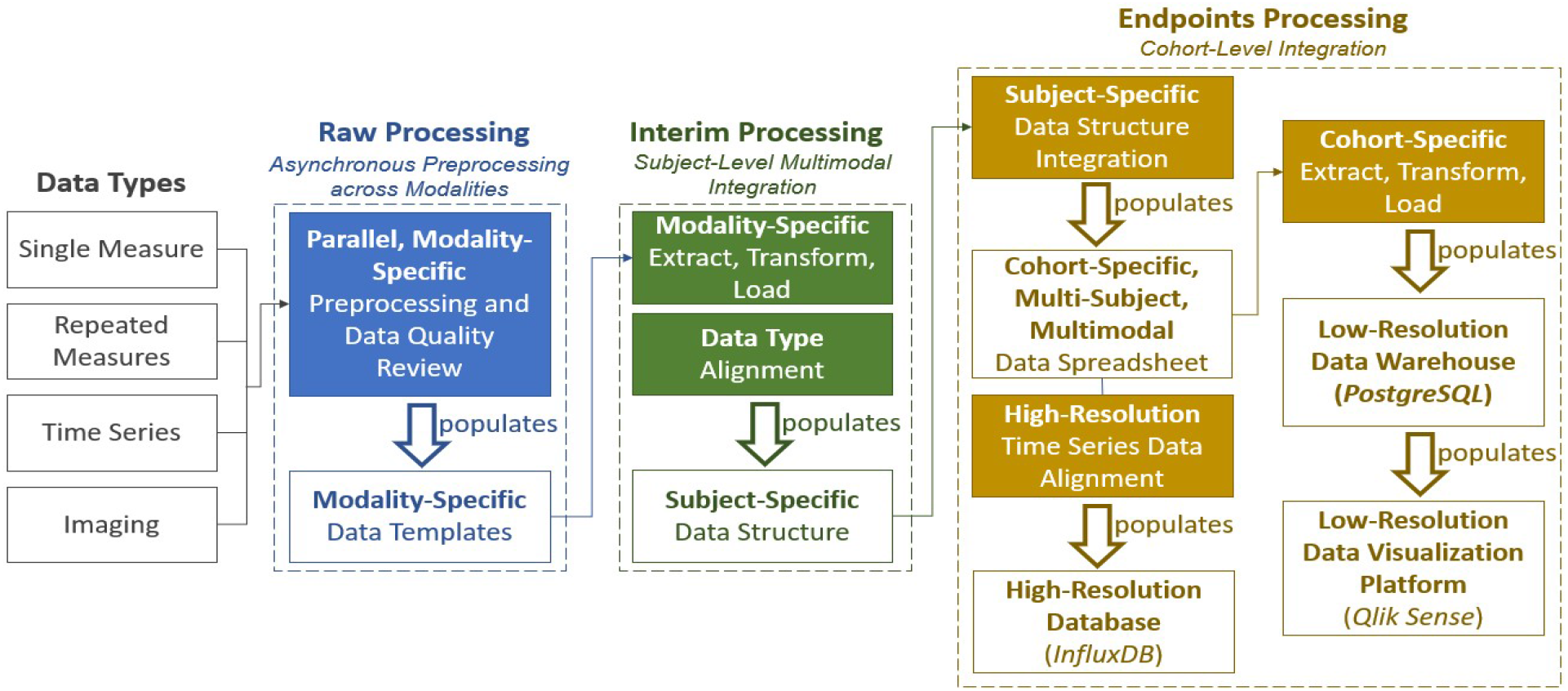
Processing Stage Flowchart: This flow chart demonstrates the functionality of each of the three data processing stages (“raw”, “interim”, “endpoints”). The primary role of each stage is visualized; modality processing during “raw”, subject-integration during “interim”, and cohort-integration during “endpoints”. Data can be traced from raw, collected data type, to the databases and data visualization platform. It is important to note that the “Imaging” data type represents the quantitative data derived from imaging data in this figure. Raw imaging data is stored in a *Flywheel* database.

### Raw Processing: Asynchronous Preprocessing across Modalities

The pipeline starts in the “raw” processing stage with preprocessing and the autonomous and manual data quality-review of disparate data types within a flexible file architecture. The “raw” processing stage is comprised of three substages. In the first substage, data collection and processing occur in dedicated modality-specific subdirectories within the “raw” directory. Distinct space allocation for each modality permits asynchronous and parallel data processing across disparate laboratories and personnel that minimizes bottlenecks. The second substage of the “raw” processing stage is data quality review.

Autonomous data quality criteria are applied and verified by manual data quality review by trained laboratory personnel. In the third substage, a standardized data template and pipeline import adapter is developed for each data modality for integration with other modalities. Each data template includes subject identification, modality-specific data elements, a time axis (if applicable), and a reference timepoint used to align the time axis to experimental periods. The pipeline import adapter is developed in collaboration with data modality experts to map quality-reviewed data into the standardized data template.

### Interim Processing: Subject-Level, Multimodal Integration

Time synchronization and intra-subject data consolidation occur in the “interim” processing stage. A subject-specific data structure is created to store all aligned and time-synchronized data modalities as the pipeline advances. This data structure includes metadata describing the animal and experimental characteristics and associated data directories. The data structure is populated from each modality’s standardized data templates (created during the “raw” processing stage). For each modality, a dedicated ETL script is developed for import into the data structure.

Time series data that have been experimentally aligned and down sampled into 15-second epochs during the “raw” processing stage are imported into the subject’s data structure. These modalities include hemodynamics and other physiological parameters, NOM, CPR performance, and EEG data. Next, single measure and repeated measures data are imported and aligned with the corresponding experimental period and respective 15-second epoch. These modalities include *REDCap*, microdialysis, mitochondrial respirometry, serum biomarkers (proteins and metabolites), and derived imaging metrics data. These integrated data are exported as a tabular .*csv* file, classified as the “summary” file to the subject’s folder in the “interim” directory, along with the subject’s data structure.

### Endpoints Processing: Cohort-Level Integration

The “endpoints” processing stage performs cohort-level multimodal data integration and incorporates the “extract-transform-load” (ETL) data pipeline into the *PostgreSQL (PostgreSQL Global Development Group, Berkeley, California, USA)* data warehouse for *Qlik Sense (Qlik, King of Prussia, PA, USA)* dashboard visualization, and into the *InfluxDB (InfluxData Inc., San Francisco, CA, USA)* high-resolution database for downstream machine learning applications. High-quality, multimodal subject data are compiled into a single data sheet for each cohort. The data sheet is two-dimensional with columns containing all measured data elements as well as identifiers for subject, experimental period, and period-specific time, and rows containing incremental data samples. This data sheet is populated from the individual subjects’ tabular .csv files located in the subject’s folder in the “interim” directory and saved in the subject’s folder in the “endpoints” directory where it is queried for further data warehousing.

### Extract-Transform-Load (ETL) Pipeline

The cohort-specific, multi-subject, multimodal data spreadsheets are used to populate the high-resolution database (*InfluxDB*) and the low-resolution data warehouse (*PostgreSQL*) which ultimately populate the *Qlik Sense* dashboard. Methodology and architecture related to high-resolution data preparation, the cohort-specific ETL pipeline, and the data visualization dashboard (*Qlik Sense*) design are presented below.

#### High-Resolution Data Preparation and Import

An *InfluxDB* database was created for storage of high-resolution (100 Hz) time series data (F.M., L.S., F.T.) and to support preclinical predictive model development. Metadata saved during manual quality-review of time series waveforms are applied to high-resolution time series data. Single measure and repeated measures data are then time aligned with the quality-reviewed high-resolution time series data. Cohort-level integration is performed, and high-resolution multimodal data *.csv* spreadsheets are generated for import into the *InfluxDB* database.

#### Low-Resolution Data Warehousing

An ETL-based data warehousing approach is used to integrate data from multiple experimental models into a single, standardized data warehouse (Fig. 3). Here, the ETL is implemented in *Python (Python Software Foundation, Wilmington, DE, USA)* and loaded into a *PostgreSQL* data warehouse. The data warehouse compiles data from *REDCap* instruments, manifest files, summary files, and the *Freezerworks* database. These data are stored in tables that are related using the unique identifier of each subject. This enables the search and modification of each data modality separately and facilitates dynamic and selective loading into the *Qlik Sense* dashboard.

**Fig. 3.**
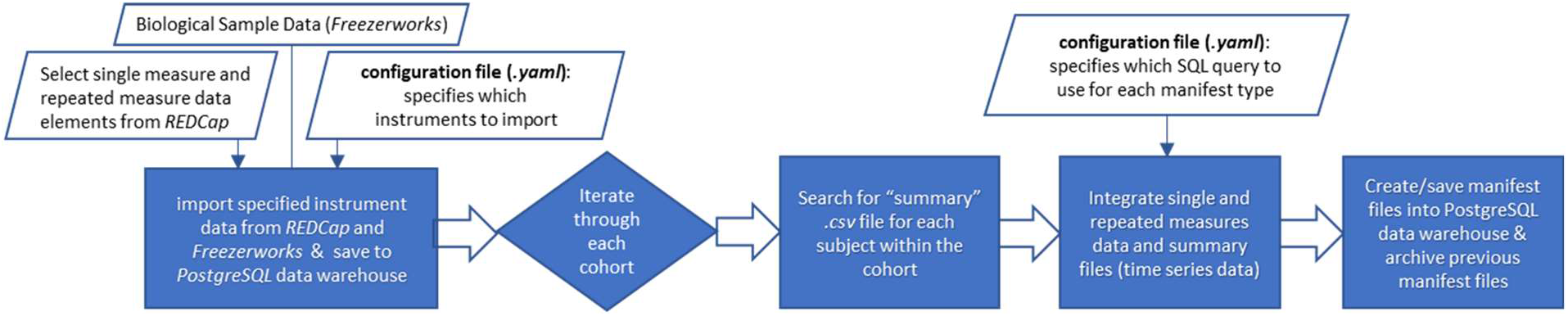
Cohort-Specific Extract, Transform, Load (ETL) Flowchart: This flowchart demonstrates the functionality of the ETL script used for manifest file creation and low-resolution data warehouse (*PostgreSQL*) data population. Configuration files specify the *SQL* queries to be used for each data type’s manifest as well as which instruments should be integrated from *REDCap* and *Freezerworks* for each cohort. The script iterates through each cohort, then each subject, searching for its “summary” data file. This file’s data is then combined with the single and repeated measures data pulled from *REDCap* and *Freezerworks* to generate manifest files that are formatted for, and saved to, the *PostgreSQL* data warehouse. All previous manifest files are stored as backup files for redundancy and version tracking in the file structure.

#### REDCap Integration

The ETL pipeline extracts data from *REDCap* using customized application programming interface (API) requests. Relevant *REDCap* instruments were identified and include “Demographics”, “Monitoring”, “Tissue Collection”, “Resuscitation”, and “Baseline” single measure instruments and “Blood Gasses”, “Cardiac Outputs”, and “ECMO” repeated measures instruments. A configuration file containing this list of instruments was used to extract instrument-specific data from *REDCap* into a dedicated table within the *PostgreSQL* data warehouse.

*REDCap* fields containing multiple choices are concatenated into a single field. Repeat instances are concatenated for ease of storage. Additional transformations are performed, including the creation of tables to associate the *REDCap* “Arm” names (e.g., cohort names) with the subjects within the cohort, the extraction of the experimental period names from each cohort, and the creation of one master “group” column that combines the various experimental groups (e.g., “sham”, “control”) from each cohort. These transformed data are then loaded into the *PostgreSQL* data warehouse and designated tables for each *REDCap* instrument are created.

#### Freezerworks Integration

The ETL pipeline extracts data from *Freezerworks* using *Freezerwork’s* API request and the unique subject identifiers in each cohort from the associated “interim” folders. Information pertaining to each subject’s biological sample collection is combined into a summary file for each cohort and then into a “master” summary file containing all the cohort’s summary files for upload into the *PostgreSQL* data warehouse.

#### Manifest Generation

Manifests are tabular records that summarize data modality availability and processing status for each subject across a cohort. Manifests are created by applying logical transformations to inputs including the summary “endpoints” data sheets (see “Endpoints Processing”) and modality collection checklists that are inputted into *REDCap* during the experiment. The output is a categorical variable with the following allowable values and corresponding definitions: −1, data collection not marked complete; 0, data not collected; 1, data collected, but not quality-reviewed; and 2, data collected and quality-reviewed.

Logic for identifying the presence of data and determining data availability and processing status are saved in *SQL* queries. These *SQL* queries are then run in *PostgreSQL* using a *Python* script to populate the tabular “manifest”, which is then exported as a *.csv* file in the “manifest” file directory and saved in the *PostgreSQL* data warehouse in a dedicated table. Exception logic in the *Python* script ensures that errors in the data, missing data, and incorrect column naming are identified and recorded without causing an abortion of the script’s run.

### Workflow Automation

The migration of processed data into the *PostgreSQL* data warehouse is automated to ensure updated data access and an accurate reflection of data availability and processing status.

The ETL *Python* script was developed and version-controlled using the *GitHub (Microsoft Corporation, Redmond, WA, USA)* repository. *GitHub* is a widely used online software development platform that enables storage, tracking, collaborative development, and publishing of software projects. This ETL script is packaged into a single, deployable, executable using *Docker (Docker, Inc., Palo Alto, CA, USA)*. *Docker* is a platform for the creation and management of containers. Containers are packaged executables that contain source code, in this case, the ETL *Python* script, and operating system dependencies that are required to run the source code in any environment. A *Dockerfile*, which serves as the guidelines to build a *Docker* image, was developed, and a *Docker* image was subsequently generated.

A *Docker* container, which is a running instance of the *Docker* image is executed daily by *Kubernetes (Linux Foundation, San Francisco, CA, USA)*. *Kubernetes* is a container-orchestration tool that facilitates the management of the *Docker* container.

The *GitHub* directory containing the ETL *Python* script and *Docker* container specification script is monitored by *Red Hat Quay (Red Hat Software, Raleigh, NC, USA)* for any modifications. *Red Hat Quay* is a container registry that interprets the container specification script and builds, analyzes, and distributes *Docker* images. When code changes are committed to *GitHub*, *Red Hat Quay* automatically detects the commit and triggers the build of a new *Docker* image.

Once *Red Hat Quay* generates a *Docker* image, an “instance” of this container is scheduled to run daily on a *Kubernetes* cluster. A .*json* configuration file is imported into *Kubernetes*, which specifies the execution schedule, the location of the image, and the minimum central processing unit (CPU) and memory requirements, among other settings. *Kubernetes* subsequently automates the execution of the ETL script to ensure that the *PostgreSQL* data warehouse is updated daily. The steps of the workflow automation pipeline are demonstrated in Fig. 4.

**Fig. 4.**
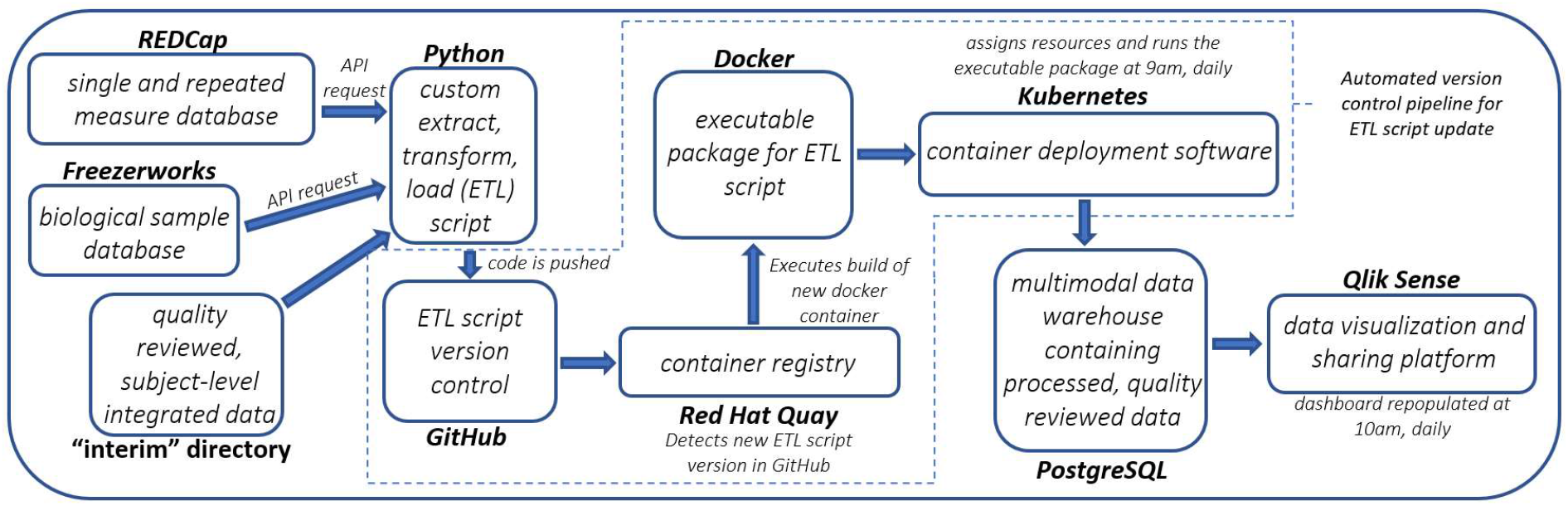
Workflow Automation Pipeline: This pipeline illustrates how components of the data framework are automatically updated when changes are made to the ETL script and how the *PostgreSQL* data warehouse and *Qlik Sense* database are automatically updated every day. Changes made to the ETL *Python* script are pushed to *GitHub* and detected via *Red Hat Quay,* triggering a rebuild of the *Docker* container. *Kubernetes* deploys the *Docker* container daily which causes the *PostgreSQL* data warehouse to be updated. The *Qlik Sense* dashboard is also updated daily.

### Data Visualization Dashboard

Compiled data streams within the *PostgreSQL* data warehouse are imported into *Qlik Sense*, a data discovery, visualization and export tool. *Qlik Sense Enterprise* is a commercial web applet that has been made accessible within our institutional network. Customized data visualization dashboards were developed to empower users to explore relevant selections of data, quickly visualize cross-sectional, time series data and perform summary analytics.

*PostgreSQL* database credentials were provided to *Qlik Sense* to permit direct query and import of data tables. Specifying table associations within *Qlik Sense* results in the joining of tables to create a single consolidated table with all unique data fields that may be ubiquitously queried by the subject identifier. Table associations and identifiers are visually depicted in Fig. 5. A built-in scheduling tool within *Qlik Sense* performs a daily query of the data warehouse and refreshes the consolidated data table.

**Fig. 5.**
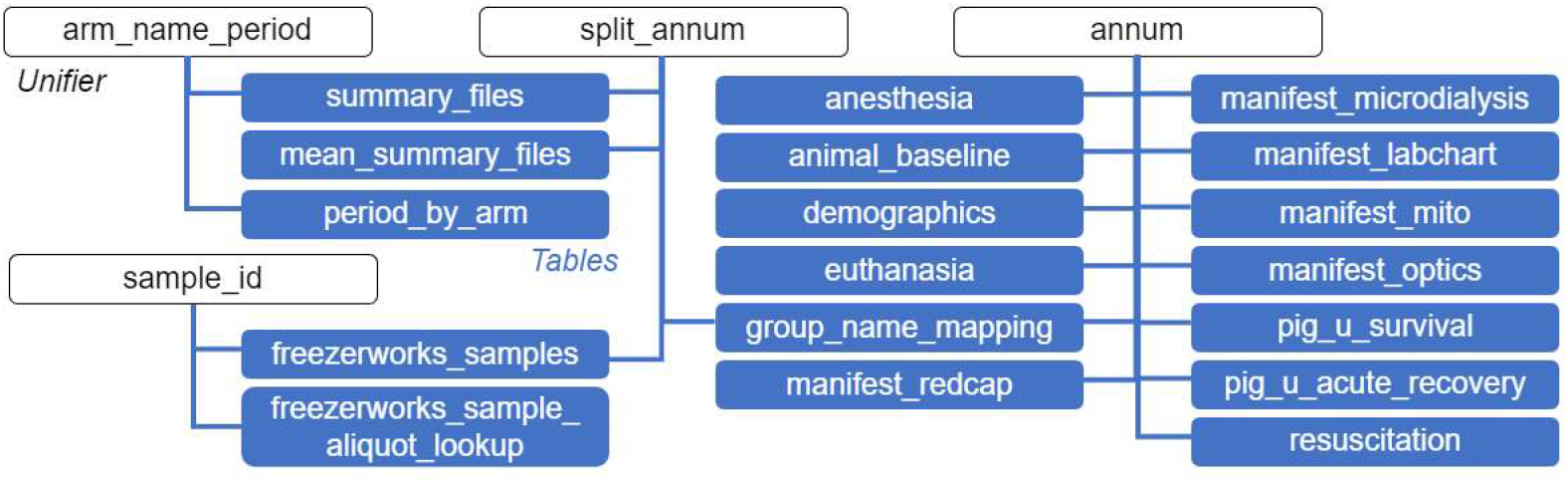
*Qlik Sense* Dashboard Table Associations: This diagram displays all of the associated tables imported into *Qlik Sense* and the “unifiers” used to associate them (“arm_name_period”, “split_annum”, “annum”, “sample_id”). “annum” is a subject-level identifier comprised of the experiment date in the format “YYMMDD” and the number assigned to the animal by the vendor. Similarly, “sample_id” is a sample-level identifier that is comprised of the date of biological sample collection in the format “YYMMDD” and the number assigned to the animal by the vendor. “split_annum” is a subject-level identifier that is only comprised of the number assigned to the animal by the vendor. This is used to link data from *Freezerworks* to data from other sources because the date of biological sample collection, which is tied to *Freezerworks* data, is not always the same as the experiment date. “arm_name_period” is a cohort-level identifier comprised of the CHOP IACUC protocol number, the year of the first experiment, a unique four-character alphanumeric cohort identifier, and the principal investigator’s last name (e.g., 1261 2022 NOME Ko).

The design of the *Qlik Sense* dashboard, including featured data fields and layout, was iteratively proposed and reviewed by research team members to ensure broad utility. The final dashboard features a landing page that centralizes data selection filters, graphical display pages that summarize selected data, a tabular page depicting the status of data processing within selected subjects, and a summary data table that permits export of all selected data. Images and further explanations of the dashboard pages are included in the “Supplementary Materials” section.

### Personnel

To develop and manage a data framework such as the one described presently, several key personnel roles are needed. First, a bioinformatician oversees data entry, the addition of new data types and cohorts, and designs new front-end data acquisition systems. This role serves as the liaison between the experimental team and the data integrationist. A data integrationist is needed to architect an automatically updated database, manage all API requests, and develop a custom ETL pipeline. Finally, a project manager guides the framework, communicating directly with principal investigators to ensure that the data needs of the lab are being met. This person will ensure that the data framework is designed to answer the research questions of interest. To maintain objectivity and avoid bias, the member of the bioinformatics team that performs data quality-review (the only qualitative portion of the data framework) is always blinded to the experimental group of the subject they are reviewing

## Results

An important contribution of these pipelines and interactive databases is the ability to generate data-driven answers that elucidate underlying physiology, injury mechanisms, and potential therapeutic targets. Here, we present a use case of the data framework to uncover intra-arrest predictors of resuscitation success. We specifically focus on the application of the novel data framework that has not been previously reported and augment the methodology included in prior work (45).

### Intra-Arrest Prediction of Resuscitation Success: A Use Case

Pediatric cardiac arrest has a high burden of mortality and neurological morbidity (46). Over the last five years, the RSC has performed ∼400 prospective cardiac arrest studies. A subset of these subjects has been included in this use case. Specifically, this study included data collected in 1- to 2-month-old swine (*sus scrofa domesticus*) that underwent either a sepsis-associated or asphyxia-associated cardiac arrest followed by 10-25 minutes of CPR (Fig. 6).

**Fig. 6.**
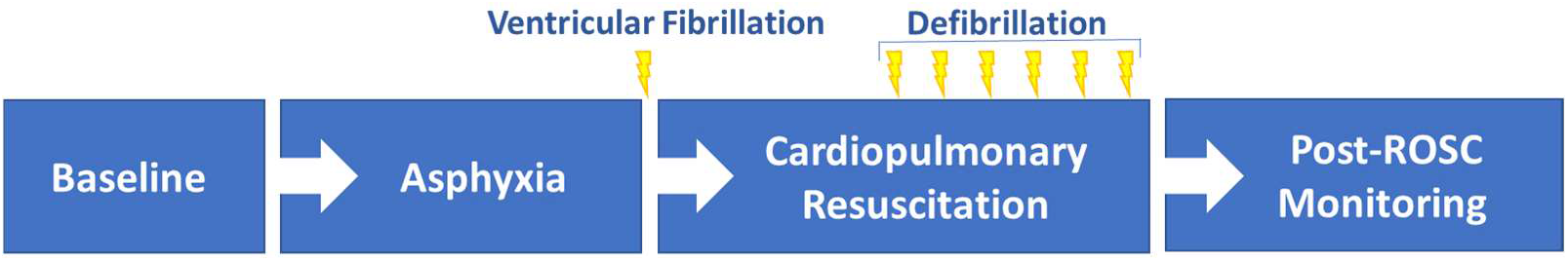
Large Animal Cardiac Arrest Model: The large animal experimental protocol for cardiac arrest and cardiopulmonary resuscitation injury model is shown here. Data collected within or across experimental periods is saved with metadata that enables association with an experimental period and timestamp. Associating all single measure, repeated measures, time series, and imaging data with an experimental period and timestamp allows data to be organized chronologically and within context of the experiment.

The objective of this study was to identify the optimal model for the prediction of return of spontaneous circulation (ROSC) using logistic regression analysis of low-resolution (15-second epoch) time series physiological data sets collected during the first 10 minutes of CPR. Utilization of the data framework to achieve this goal is demonstrated by the steps below.

### Raw Data Processing

Physiological waveforms were recorded in *LabChart*. These waveforms included AoP, RaP, PaP, electrocardiography (EKG), peripheral oxygen saturation (SpO2), capnography, and derived measures including systolic and diastolic blood pressures, heart rate, and EtCO2. *LabChart* data files (.*adicht*) for each subject were segmented by experimental period (e.g., “baseline”, “asphyxia”, “cpr”, “postROSC”; see Fig. 6), down sampled by block-averaging from a raw acquisition rate of 1000 Hz to 100 Hz and exported as a .*txt* file. The exported file was imported into *MATLAB* using a custom script. These data were then averaged into 15-second epochs and exported to a .*csv* file for manual data quality-review.

### Data Quality-Review

Each subject’s *.csv* file was manually quality-reviewed by a member of the bioinformatics team. During this process, baseline data were assessed to ensure normative physiological values. Ranges for normative values were derived from literature and typically reflected the 5th and 95th percentiles for low and high cutoffs, respectively. Waveforms that exceeded physiological limits at baseline were omitted for all experimental periods. Data from subsequent experimental periods were assessed for expected physiological trends (e.g., decreasing SpO2 during asphyxia, loss of blood pressure during ventricular fibrillation), to ensure that the experimental protocol was properly performed and that physiological waveforms were quantitatively accurate. If a waveform fell within physiological limits at baseline but did not exhibit expected physiologic trends, the waveform was omitted for all experimental periods. This process effectively removed measurement artifacts and poor-quality data associated with incidental failure of monitoring devices and deviations from experimental protocols. Following quality-review, the remaining high-quality data for each subject were saved as a separate .csv using a filename that conveyed that data quality-review was complete.

Inclusion and exclusion criteria, normative ranges, and mean and standard deviation data for historical time series physiological variables collected in 8-12 kg animals can be found in Table I (22,41,47–56). The average and standard deviation values presented in this table were acquired using the time series average graphing functionality of the *Qlik Sense* dashboard on the “Time Series Graphs” page using the data set analyzed in the presented use-case. The values were obtained from the last epoch (15 seconds) of the baseline period and the last epoch (15 seconds) of the asphyxia period.

**TABLE I.**
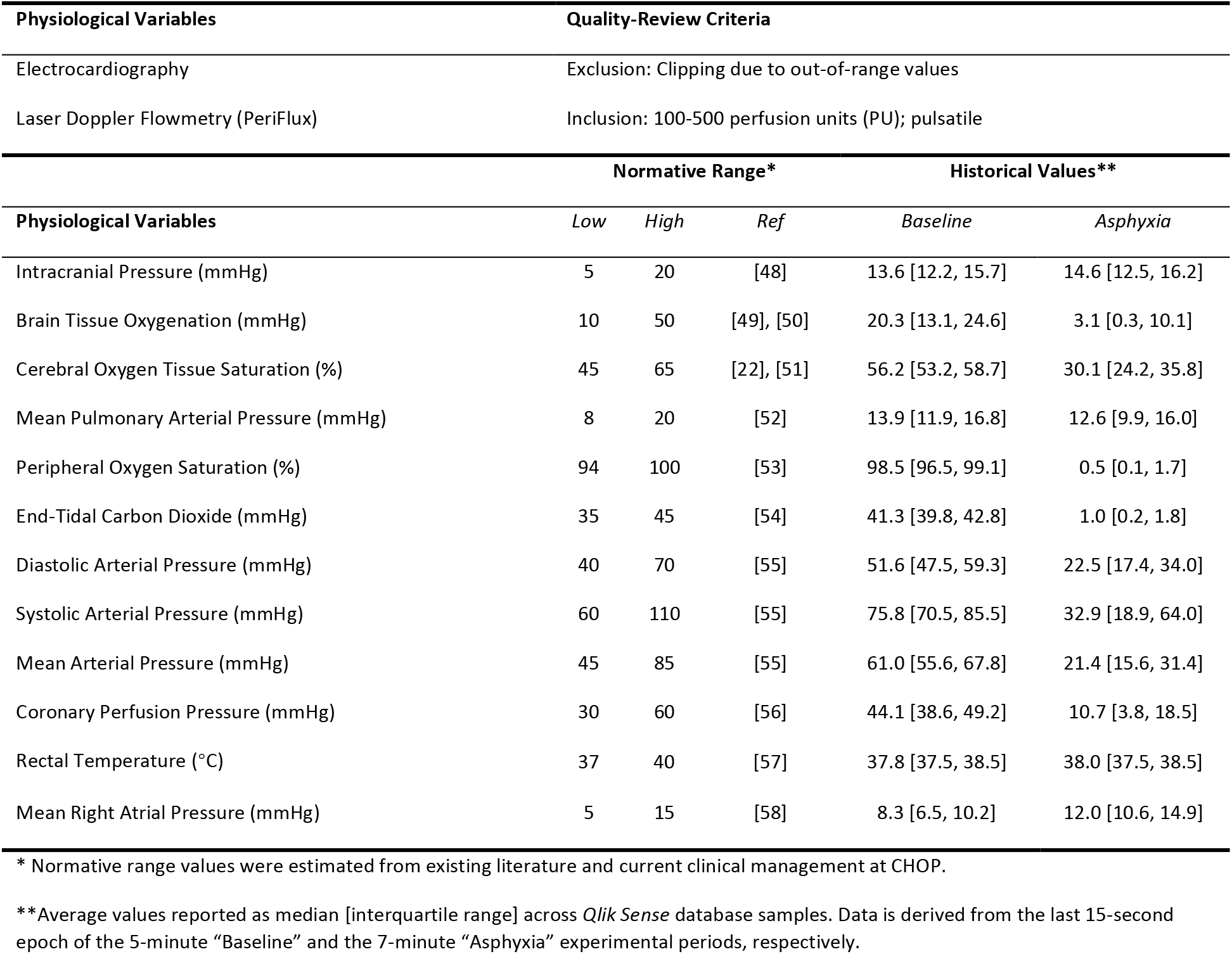
Data Quality-Review Criteria, Normative Ranges, and Average Values in Pediatric Cardiac Arrest Swine Cohort.

### Interim Data Processing

The quality-reviewed time series data .*csv* file for each subject was then read into *MATLAB*, and poor data quality omissions were propagated to the high-resolution (100 Hz) data by removing the entirety of the omitted 15-second epoch’s data. The physiological waveform data were then time-aligned and integrated with other data types (e.g., single measure, repeated measures), and the consolidated data set was organized by experimental period and stored in a *MATLAB* data structure. From this data structure, a tabular .*csv* file containing all integrated data streams was generated and saved to the subject’s “interim” folder.

### Endpoints Data Processing

The custom extract, transform, load *Python* script was executed daily by *Kubernetes*. This script scraped each subject’s “interim” folder, extracted the 15-second epoch physiological waveform data from the summary .*csv* files, compiled these data across each experimental cohort, and imported these data into the *PostgreSQL* data warehouse. The *Qlik Sense* scheduler then performed daily queries of the *PostgreSQL* data warehouse to refresh and rejoin data tables according to the user-specified table associations.

### End-User Data Extraction and Visualization

The custom *Qlik Sense* dashboard was used to compile the data set of subjects based on pre-determined inclusion and exclusion criteria, specifically, subjects who were in experimental cohorts that underwent an asphyxia-associated or sepsis-associated cardiac arrest followed by 10-25 minutes of CPR. Sham subjects that did not receive an injury were excluded.

The data set for this study was rapidly extracted using the filter functionality on the landing page of the *Qlik Sense* dashboard (Fig. 7). The landing page permits user selection of data field values that are commonly used to identify data subsets (e.g., cohort identifier, experimental model). Additionally, convenient summary statistics such as a list of subjects selected, number of subjects selected, and the range of experiment dates selected, are displayed. The “Experimental Type” was selected as “CardiacArrest”; only experimental “Cohorts” that underwent asphyxia as their arrest etiology were selected. Then, all cohort-specific groups that did not receive CPR were removed (e.g., “Sham”). This yielded a data set of all subjects who underwent an asphyxia-associated cardiac arrest and subsequent CPR. Detailed steps for visualization of the selected data in the *Qlik Sense* dashboard set are expanded upon below.

**Fig. 7.**
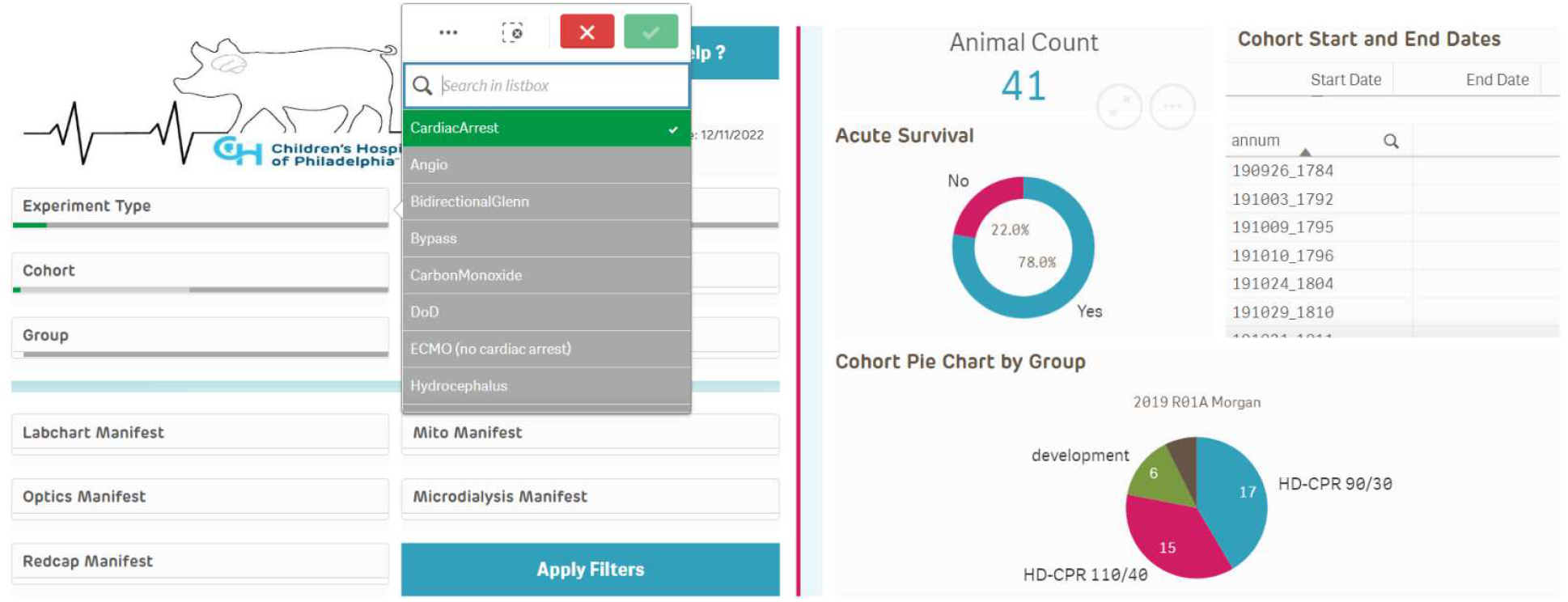
Landing Page of the *Qlik Sense* Dashboard: The landing page of the *Qlik Sense* dashboard allows the user to perform filtering to generate a sub data set from all available data within the framework. The data filters selected on this page translate through all subsequent pages of the dashboard. The left-hand side of the page features filters at the experimental model, cohort, and subject level, as well as “manifest” filtering, which enables filtering at a data type level. The right-hand side of the page visualizes the user’s selected data set using the total “Animal Count”, a pie chart displaying the experimental groups and number of subjects, and a list of the subject IDs within the selected data set.

### Use Case Data Visualization

Calculated statistical values from the selected data set were viewed on the “Summary Parameters” page. On the “Time Series Graphing” page, the distribution of individual time series physiological waveforms during baseline, asphyxia, and CPR experimental periods were viewed, as demonstrated in Fig. S.3 and Fig. S.4 (see Supplementary Materials). Individual waveforms that were apparent outliers from the data set were carefully examined for data quality and quantitative accuracy. Additional “Average Graphs” and “Box Plot” graphing tabs enabled rapid summary of time series across the data set. For example, this enabled quick summarization of the effects of asphyxia on mean arterial pressure (MAP). At baseline, MAP (mmHg) has a mean value and standard deviation of 63.0 and 9.4, respectively; at the end of seven minutes of asphyxia, this drops to 24.9 (13.7) mmHg.

The “Split Groups” page enabled the visual comparison of time series data of two separate experimental groups or outcome groups (e.g., ROSC vs. no ROSC) in parallel as demonstrated in Fig. S.5. In this use-case, this functionality was used to compare subjects that achieved ROSC, and those who did not to identify potential predictive variables.

The “Correlation Graphing” page enabled the graphing of the correlation between two variables during the selected experimental time periods, as demonstrated in Fig. S.6. This visualization helps to identify potential physiological associations and causal relationships. The generated correlation graph displays bins of datapoints; high-density bins (which contain more datapoints) are shaded darker. The incidence is printed on each point.

Finally, the filtered data set was downloaded from the “Summary Data” page into a .csv file. In this file, every single measure, repeated measures, and time series data point is associated with an experimental group and subject ID.

### Predictive Modeling

This use case evaluated the performance of features extracted from physiological waveforms in different time segments, separately and combined, to predict ROSC during CPR. Predictive model development has been previously detailed (45) and will be briefly summarized here. This use case utilized a logistic regression model with Least Absolute Shrinkage and Selection Operator (LASSO) regularization (L1 penalty). Four additional models (logistic regression without LASSO, XGBoost, random forest and multilayer perceptron) were trained, and their performances were compared with those of logistic regression with LASSO. Because the data set was class-imbalanced (72 ROSC and 17 NO ROSC), assigned training weights were inversely proportional to class frequency. The training iteration was 100, and a scheme of nested cross-validation was used to assess model performance.

### Use Case Results

The general conclusions of the published use-case study were that prediction performances generally improved after the 6th minute of CPR, and EtCO2 yielded the best individual variable performance as a predictor of ROSC from the 2nd to the 4th minute of CPR, making it a candidate for early CPR-stage prediction. Overall, the study serves as an important foundation for improving the likelihood of achieving ROSC during CPR based on multimodal waveform feature extraction (45).

## Discussion

Preclinical, large animal models enable the *in vivo* study of neurological injury and disease with unique benefits for clinical translation. Elucidation of injury pathways facilitates the development of novel therapeutics. At present, storage architecture and integration of diverse and complex data sources poses a significant challenge to data analysis that provides access to these insights. To address this challenge, we have developed and described a custom data framework for the rapid, modular handling of heterogeneous and longitudinal preclinical research data types. To the best of our knowledge, no existing publicly available data repositories or processing platforms have developed standardized data architecture to integrate these complex data types. This work is important because it provides a generalizable framework that may be readily adopted for this purpose. These standardized, integrated data sets will facilitate data analysis as well as data dissemination. The latter is particularly relevant given the recent data accessibility mandate by the National Institutes of Health (https://grants.nih.gov/grants/guide/notice-files/NOT-OD-21-013.html). By addressing a significant hurdle in scientific discovery, these bioinformatics developments strengthen the bridge between preclinical research and clinical translation.

The presented data framework was tailored to address aspects of preclinical research data management that are distinct from clinical research. First, standardized injury models and study designs result in repeatable experimental periods that were leveraged to structure data and facilitate key cross-sectional comparisons across cohorts. New cohorts using the same injury model were easily integrated. It is important to note that comparability between study designs relied on the implementation of generalizable nomenclature to describe experimental periods and interventions such as “baseline”, “injury”, “exposure”, and “recovery”. Additionally, in preclinical research, a multitude of physiological parameters may be simultaneously collected. Within these parameters, the use of novel and/or experimental measurement parameters that do not yet have standardized data formats and are sampled irregularly are common. The framework’s period-based architecture flexibly accommodates any number of data types and only requires a variable name, timepoint, and value. This simplifies the addition of novel or derived data types. In the framework’s implementation, the harmonization of variable names and data formats across study designs permitted rapid data extraction of comparable data elements across disparate cohorts via the data dashboard. By providing a framework for the integration of preclinical data, this work aids in the preservation and dissemination of preclinical data sets that may inform future studies and would otherwise be inaccessible.

A significant advantage of the presented data framework is its adaptability. Pre-existing data framework elements enable the rapid standardization of data collection for new experimental cohorts. These standardized data are automatically integrated and made accessible within the framework. The significant reduction in time and resources necessary for data organization has driven adoption across the multi-disciplinary research center.

In utilizing the data framework for the use case, major strengths and weaknesses were realized. First, there was no easy way for users to communicate issues they encountered with the data or while using the dashboard. This prompted the addition of a help button and feedback feature within *Qlik Sense*. This enables users to suggest new features and alternate designs and to report errors within the platform.

The bioinformatics team is automatically notified when the user submits their entry. Aside from the inability to communicate shortcomings or errors within the framework, the first comprehensive use of the framework was successful. The user was able to download all data associated with the subjects of interest from one location, after tailoring the filtering criteria to the study parameters. Additionally, because the study was being performed concurrently with ongoing swine cardiac arrest experiments, new eligible data were consistently imported into the database. The automated updating of the dashboard and standardized format allowed these new data sets to be easily downloaded and integrated into the study.

Expansive efforts are ongoing to address multi-factorial challenges related to developing robust data frameworks for preclinical studies that generate large, heterogeneous data sets. The Brain Imaging Data Structure (BIDS) (57–60) addresses standardized data formats for imaging and associated metadata, and has gained increasing use in clinical data architectures and is also compatible with the *Flywheel* imaging database. The BIDS data format supports self-reported *REDCap* data, neuroimaging data (MRI and simultaneous EEG-fMRI), collected biospecimen information (*Freezerworks*), single measure actigraphy data collected daily via *Fitbit (Fitbit, San Francisco, California)* wearable sensors, and behavioral testing data collected via *NeuroMap (Delsys, Natick, MA)*. At the Laureate Institute for Brain Research, a research team has developed a clinical framework using NIH common data elements, BIDS image standardization, and scalable data management and analytics workflows that facilitate data processing, analysis, and sharing (61). Preclinical BIDS formats are in development (62); a recent survey summarizes the ongoing challenges of data standardization in preclinical imaging (63). In the current framework, we utilize *Flywheel* for raw imaging storage, however standardization for downstream image processing is ongoing and remains limited to quantitative end point data values.

The NIH Common Data Elements Repository (https://cde.nlm.nih.gov/) seeks to standardize clinical data reporting, but preclinical CDE standardization remains limited to animal models of TBI (64). Preclinical CDEs have spurred the development of data sharing platforms for preclinical TBI research that enabled elucidation of inflammatory patterns by combining data sets across multiple studies (n=1250 subjects) (29). The continued development of inter-operable preclinical CDE libraries and standardized data formatting structures is integral to maximizing data reuse and enables large-scale data-driven scientific discovery. At present, our data framework utilizes data element definitions specific to our institution. Efforts are ongoing to harmonize these definitions with existing preclinical CDEs.

Uniquely, our data framework is built to accommodate multimodal data from diverse experimental models and cohorts. Developed integration pipelines had to unify descriptors of experimental timing and protocol features to facilitate cross-sectional analysis. In addition to contextualized multimodal data integration, our research center required an interactive data visualization platform for exploratory analysis and data sharing. To the best of our knowledge, the presented framework is the first of its kind in achieving these goals.

Additional ongoing efforts include the integration of new complex and large data types. Currently, architecture exists to house and integrate quantitative data yielded from imaging data types, however, additional integration architecture must be developed to automatically link a subject’s data set in *Qlik Sense* with imaging data stored in *Flywheel*. Also, the data visualization component of the *Qlik Sense* dashboard only includes 15-second averaged data from the down sampled 100 Hz physiological data. To monitor acute physiological responses, the visualization of the high-resolution 100 Hz data will be useful. Development of a user interface that queries the high-resolution *InfluxDB* database interface is ongoing, along with the development of an automated ETL pipeline to populate the high-resolution database (similar to that used for the low-resolution *PostgreSQL* data warehouse). Also, as additional omics data types and analysis outputs emerge, a framework must be developed for the storage, accessibility, and interpretability of these large data sets.

## Conclusions

Preclinical, translational research is paramount to the mitigation of pediatric neurological injury and disease. At present, data integration significantly impedes therapeutic development and etiological discovery. Here, we present a custom data framework that is adaptable, accessible, and automated for the handling of multimodal, longitudinal data types in the preclinical research arena. This platform is scalable and will facilitate intra- and extra-institutional collaboration to maximize utilization of preclinical data sets. Through evaluation of other existing preclinical and clinical platforms, and the application in a predictive model development use case, it has been demonstrated that this data framework can serve as a foundation for next-generation preclinical studies of pediatric neurological injury and disease. We hope that the data framework and workflow methodology presented here can serve as a blueprint for other translational laboratories.

## List of Abbreviations

TBI: traumatic brain injury

FITBIR: Federal Interagency Traumatic Brain Injury Research

CDE: common data element

ETL: extract, transform, load

RSC: Resuscitation Science Center

IACUC: Institutional Animal Care and Use Committee

CRF: case report form

USDA: United States Department of Agriculture

OXPHOS: oxidative phosphorylation

ETS: electron transport system CS citrate synthase

HIC: Human Immunology Core

AoP: aortic pressure

RAP: right atrial pressure

PAP: pulmonary artery pressure

EtCO2: end-tidal carbon dioxide

CoPP: coronary perfusion pressure

CPR: cardiopulmonary resuscitation

EEG: electroencephalography

NOM: Neurometabolic Optical Monitoring

FD-DOS: frequency-domain and broadband diffuse optical spectroscopy

b-DOS: frequency-domain and broadband diffuse optical spectroscopy

DCS: diffuse correlation spectroscopy

StO2: cerebral tissue oxygen saturation

MRI: Magnetic Resonance Imaging

SWI: susceptibility weighted imaging

DTI: diffusion tensor imaging

MRS: magnetic resonance spectroscopy

ASL: arterial spin labeling

BOLD: blood oxygenation level dependent

CEUS: contrast enhanced ultrasound

RF: radio frequency

API: application programming interface

CPU: central processing unit

ROSC: return of spontaneous circulation

MAP: mean arterial pressure

LASSO: Least Absolute Shrinkage and Selection Operator

## Declarations

### Ethics approval and consent to participate

Not applicable.

### Consent for publication

Not applicable.

### Availability of data and materials

The datasets generated and/or analyzed during the current study are not publicly available but are available from the corresponding author on reasonable request.

## Authors’ contributions

HG, VP, T. Ko, WL, T. Kilbaugh, and RM contributed to the data framework conception and design. All authors assisted with data collection. HG, VP, T. Ko, L. Silva, J. Slovis, FT, T. Kilbaugh, and RM interpreted the results. HG, VP, T. Ko, L. Silva drafted and edited the manuscript. All authors provided critical revisions of the manuscript. MH, WB, FT, T. Kilbaugh, and T. Ko provided funding for experiments that enabled development of the data framework presented here. All authors read and approved the final manuscript.

## Acknowledgements

We thank the Department of Veterinary Resources at the Children’s Hospital of Philadelphia as well as all current and former members of the large animal research team at the CHOP Resuscitation Science Center of Emphasis. Additionally, we thank the Arcus Library Science Team within the Department of Biomedical and Health Informatics at CHOP. We would also like to recognize the CHOP Research Institute Frontier team for their guidance and support, as well as the Chappell-Culpepper Foundation.

