## Supplementary Materials for "A Template for Translational Bioinformatics: Facilitating Multimodal Data Analyses in Preclinical Models of Neurological Injury"

### Qlik Sense Dashboard Visualization Features

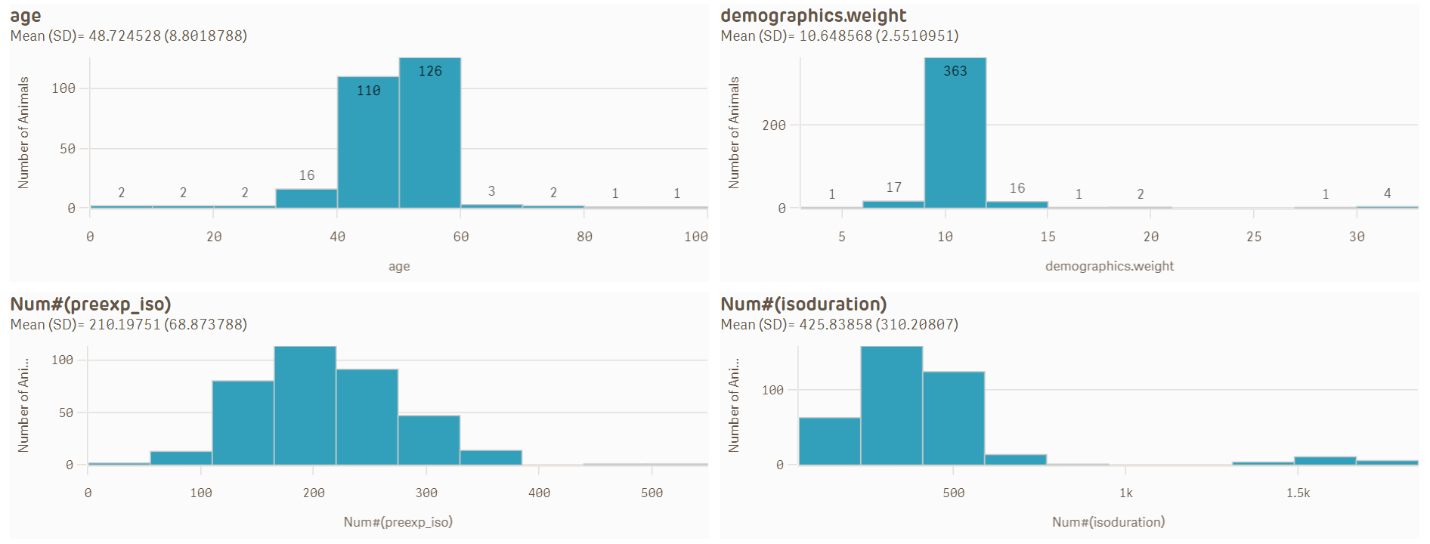

Fig. S.1. “Summary Parameters” Page of the Qlik Sense Dashboard: On the “Summary Parameters” page, bar graphs are used to visualize the distribution of specific variables within the selected data set. The “general” parameter set shows users a data set-level view of the distribution of age, weight, pre-experimental isoflurane duration, and total isoflurane duration. Different tabs contain parameter sets relevant to each experimental model. For instance, the “Cardiac Arrest” parameter set contains variables related to CPR performance and the number of shock attempts required to achieve ROSC.

A “Summary Parameters” page contains bar charts and summary values of experimental variables relevant to each experimental model (see Fig. S.1). Relevant variables were chosen for each experimental model after discussion with principal investigators and researchers. This page helps users visualize general trends related to their selected data set before conducting further exploratory analysis on the subsequent pages.

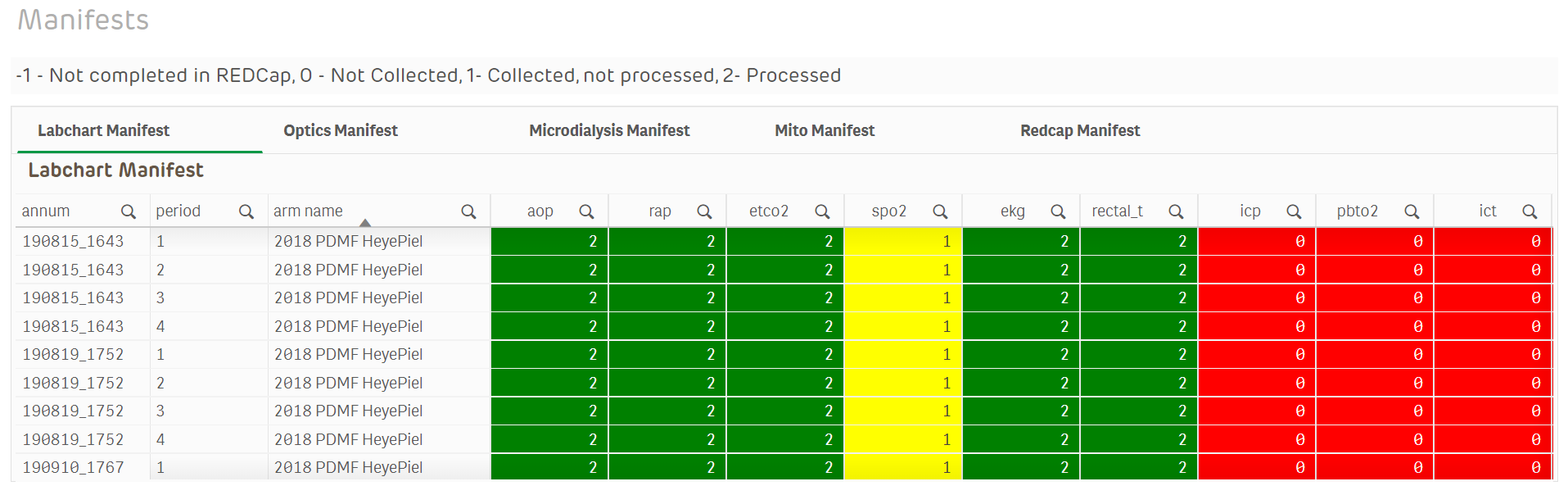

Fig. S.2. “Manifests” Page of the Qlik Sense Dashboard: The “Manifests” page displays the collection and processing status for each variable, each period, and each subject in the selected data set. A “0” (red) indicates that data was not collected, a “1” (yellow) indicates that data was collected but not processed, and a “2” (green) indicates that data was collected and processed. For example, for animal 190815_1643, oxygen saturation (“spo2”) data was collected but not processed for all four experimental periods. This page enables variable-specific filtering to remove subjects from the selected data set that do not contain quality-reviewed data for a variable of interest.

A “Manifests” page displays surrogate values that represent data type availability, quality, and usability for each subject and measurement modality across a cohort (see “Manifest Generation” and Fig. S.2).

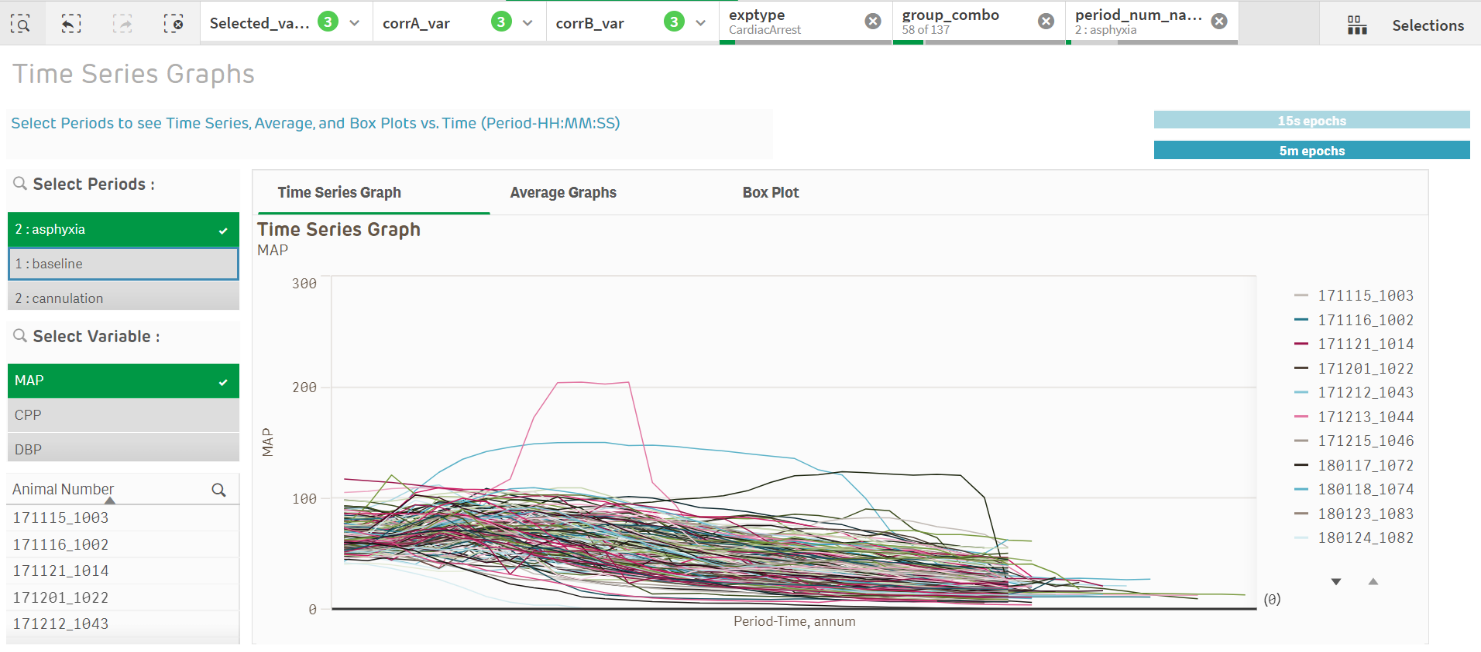

Fig. S.3. “Time Series Graphing” Page of the Qlik Sense Dashboard: The “Time Series Graphing” page facilitates plotting of the selected data over time for the selected variable and experimental period(s). Subsets of subjects can be omitted directly from this page by deselecting the subject ID in the “Animal Number” list. The “Average Graphs” tab plots the average +/- standard deviation of the selected data set. The “Box Plot” tab is explained in Fig. S.4. The user can toggle between 15-second and 5-minute discrete intervals to change sampling granularity for each of the three tabs.

A “Time Series Graphs” page enables users to visualize all the time series data associated with their filtered data set and further filter by experimental period and variable through dynamic plotting (see Fig. S.3). Users can examine the distribution of individual time series physiological waveforms during different experimental periods to find outliers or look at trends. Alternate windows on this page allow visualization of the plotted average and standard deviation, and box plots with basic statistics (median, first quartile, third quartile, box end + (1.5 x interquartile range), box end – (1.5 x interquartile range)) (see Fig. S.4). The user can also toggle between 15-second and 5-minute discrete intervals to change sampling granularity.

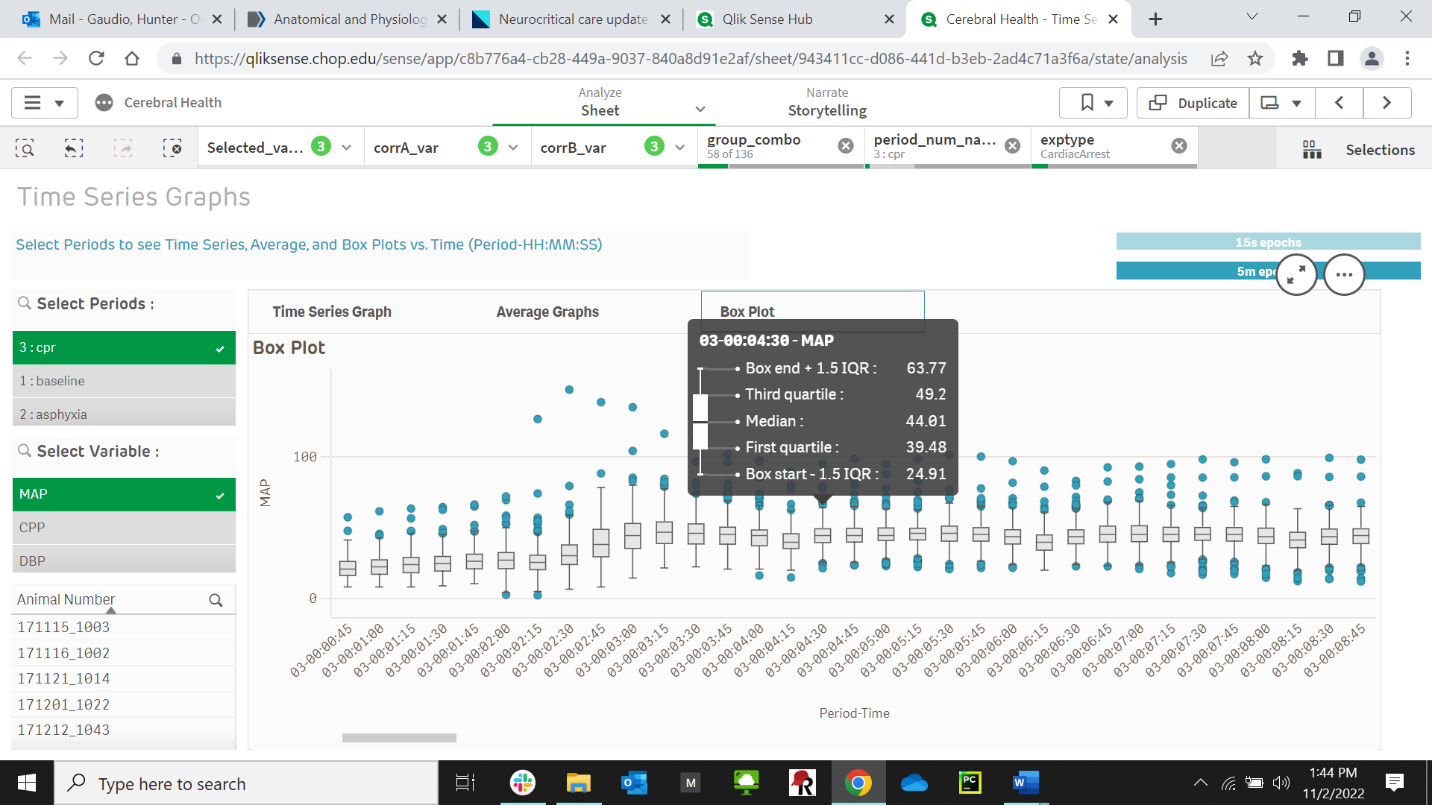

Fig. S.4. “Box Plot” Tab of the “Time Series Graphing” Page of the Qlik Sense Dashboard: The “Box Plot” tab of the “Time Series Graphing” page enables the identification of individual outlier points for each variable, over the course of each experimental period. Statistics generated for each point include median, first quartile, third quartile, box end + (1.5 x interquartile range), box end – (1.5 x interquartile range), and outliers outside of this range (blue dots).

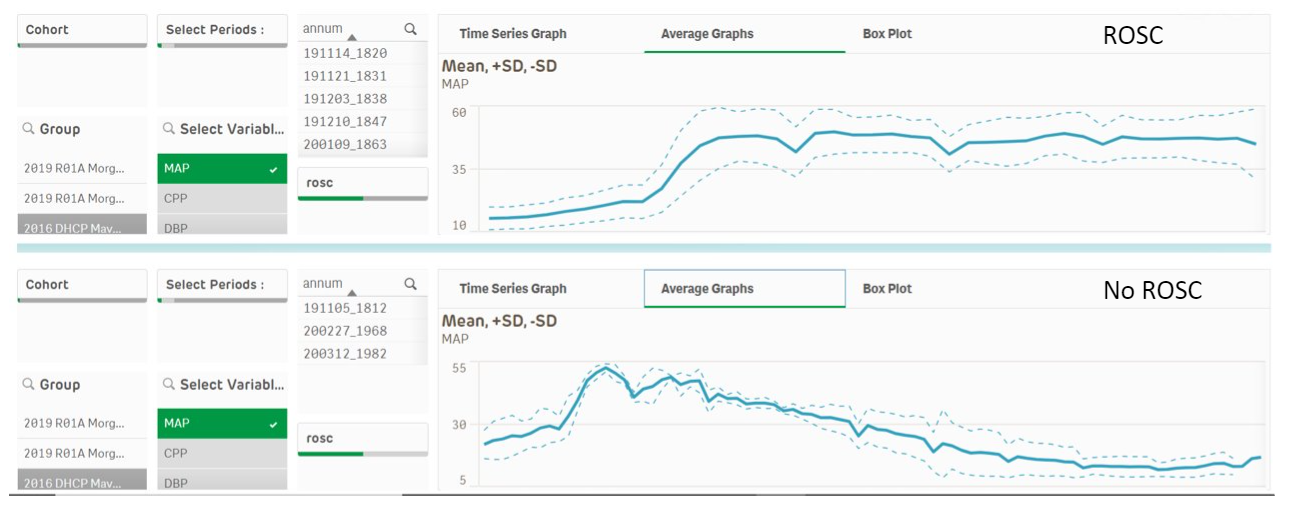

Fig. S.5. “Split Groups” Page of the Qlik Sense Dashboard: The “Split Groups” page enables parallel graphing for comparison of two data subsets. This page is used to plot either two separate data sets for the same variable, or two separate variables for the same data set. The example shown in the above figure is the evaluation of MAP during the CPR period in the subjects that achieved ROSC vs. those that did not. The “rosc” filter is used to select “1” or “0,” indicating “ROSC” or “No ROSC,” respectively. To visualize the influence of ROSC, the same cohort, period, and variable was selected in both groups.

A “Split Groups” page allows users to compare two different cohorts, or different subsets of subjects within a cohort at the same time, graphed adjacent to each other (see Fig. S.5). The design of this page presents a workaround to *Qlik Sense’s* inability to plot two separate groups on the same graph. Both plots have the same alternate windows from the “Time Series Graphs” page for average or box plot graphing.

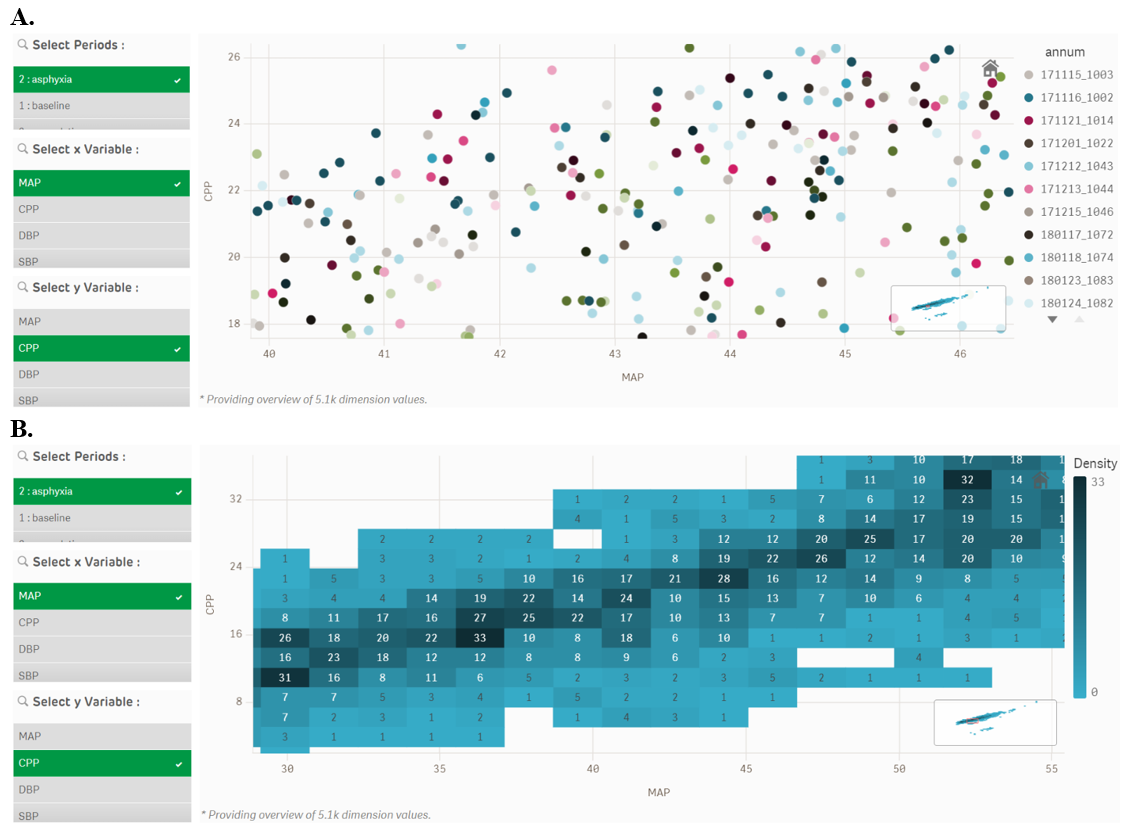

Fig. S.6. “Correlation Graphing” Page of the Qlik Sense Dashboard: The “Correlation Graphing” page enables users to visualize the relationship between two variables for the selected data set. This figure displays the relationship between mean arterial pressure (“MAP”) and coronary perfusion pressure (“CPP”) during the “asphyxia” experimental period. Depending on whether data density exceeds a size threshold predefined in Qlik Sense, data points are depicted as raw values (**A.**) or as a density plot (**B.**). (**A.**) Data sets that fall below the size threshold predefined within Qlik Sense cause individual points to be plotted; points are color-coded by subject. (**B.**) Data sets that fall above the size threshold are plotted in tiles; data density in each tile is indicated by shading and displayed incidence values.

A “Correlation Graphing” page allows the user to visualize the quantitative relationship between two variables (see Fig. S.6). Corresponding values for the selected x-axis variable and the selected y-axis variable are plotted for all subjects at all timepoints within the selected data set. For smaller data sets, individual points are color-coded by subject. Selecting a data point reveals the timepoint and subject ID corresponding to that datapoint. For larger data sets where the point density exceeds a preset threshold in the Qlik Sense dashboard, a density plot is generated to visualize the distribution of data across the x- and y-axes. Data points are binned into 2D tiles where the tile is labeled with the number of data points contained within the x- and y-axis boundaries of the tile and shaded based on the relative distribution of data points across tiles. Selecting a data tile reveals a list of subjects and timepoints whose values are contained within the tile.

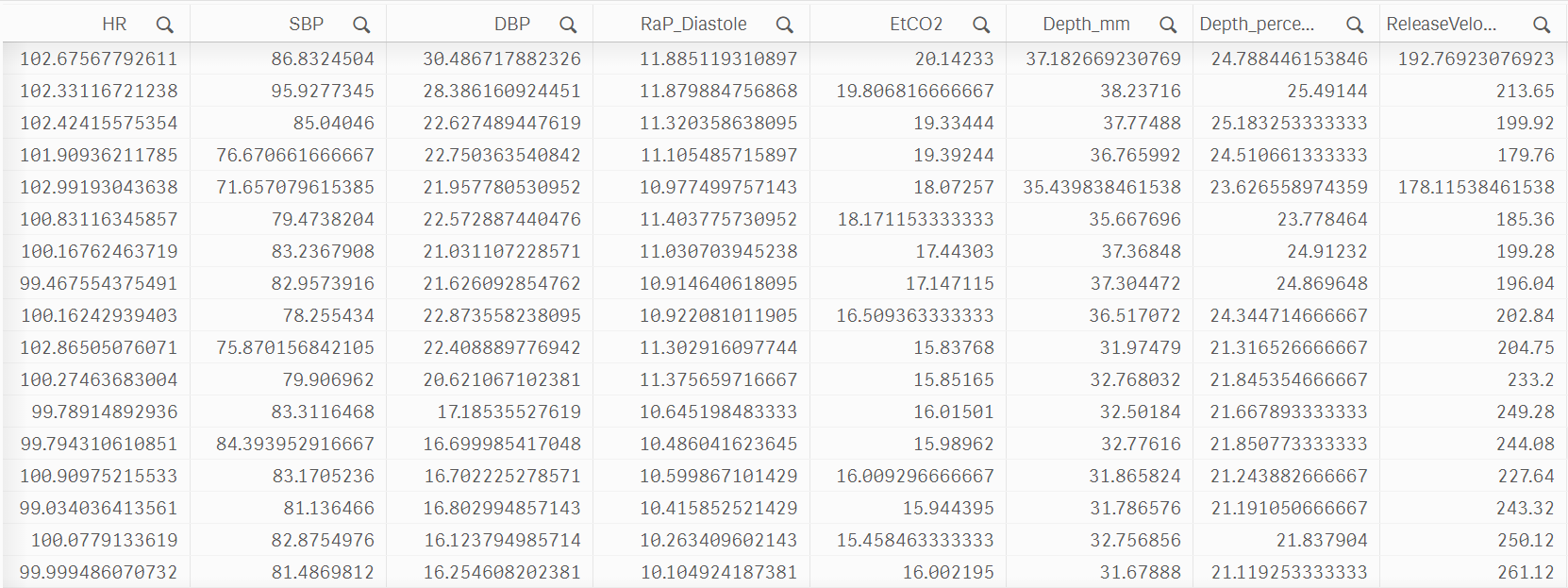

Fig. S.7. “Summary Data” Page of the Qlik Sense Dashboard: The “Summary Data” page displays all data elements for the generated data set. From this page, the full data set can be exported for further preprocessing from this sheet as an image (.png or .jpeg), .pdf file, or .csv file. Clicking the magnifying glass icons next to the variable names allows the user to sub-filter the data by range and other numerical filters. A subset of the total number of variables exported is shown. In total, a dataset containing 431 time-synchronized variables was exported for the presented use case (see “Results”).

A “Summary Data” page displays all the data in the filtered data set as a tabular sheet that can be further filtered by individual variables or exported locally as an image (.*png* or .*jpeg*), .*pdf* file, or .*csv* file for further analysis (see Fig. S.7).

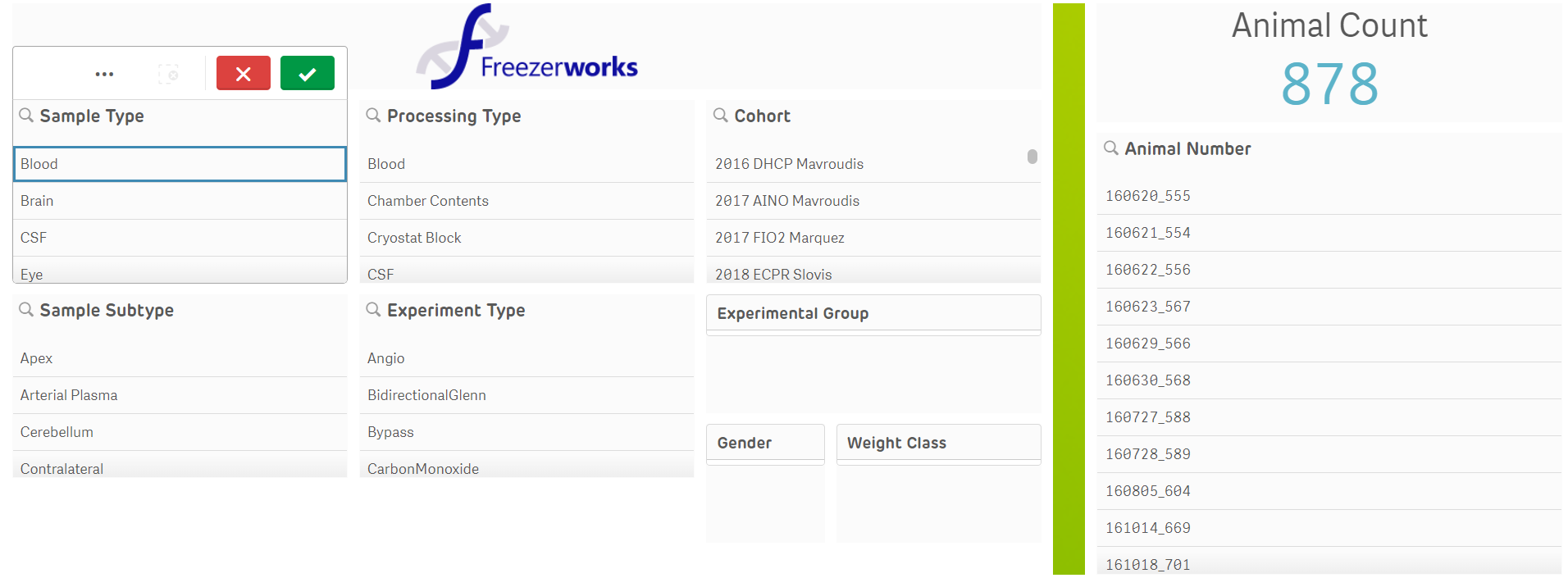

Fig. S.8. “Freezerworks Filters” Page of the Qlik Sense Dashboard: The “Freezerworks Filters” page displays all filtering criteria that pertains to the collected biological samples stored within the RSC’s biobank. From this page, filters can be applied to generate an “Animal Number” list for data download or sample request from the biobank for further downstream processing.

The “Freezerworks Filters” (see Fig. S.8.) and “Freezerworks Data” pages function identically to the landing and “Summary Data” pages, respectively, but pertain only to data pulled from the *Freezerworks* database. Filtering by variables related to biological sample collection and subsequent viewing of a tabular datasheet that groups these variables facilitates the exploration of the database with biological sample availability as the priority. This is a valuable feature for principal investigators interested in conducting retrospective studies utilizing the RSC’s extensive biobank.
